# Fusion2AI: functional re-annotation of the bacterial transparent matter proteins

**DOI:** 10.64898/2026.09.28.755017

**Authors:** Aiden Maloney-Bertelli, Yana Bromberg

**Affiliations:** Department of Computer Science, Emory University, 400 Dowman Drive, Atlanta, GA, 30322, USA; Department of Biology, Emory University, 1510 Clifton Road NE, Atlanta, GA, 30322, USA

**Keywords:** protein function annotation, Fusion, large language models, bacterial genomics, metagenomics, Gene Ontology

## Abstract

Determining the biological function of bacterial proteins remains a central challenge in microbiology. We previously developed Fusion, a reference-free scheme that clustered the tens of millions of known, unique, bacterial proteins into about four hundred thousand groups of functionally similar proteins, i.e. *functions*. For any newly identified proteins, Fusion function assignment could be performed using function-aware sequence alignment to each *function*’s representatives. However, Fusion functions lack human-readable descriptions of their functionality.

Here, we leverage large language models (LLMs) to synthesize scattered, partial annotations of functions of individual proteins into coherent English-language descriptions for *∼*100,000 clusters, i.e. a quarter of all Fusion functions. From these descriptions, we extract Gene Ontology (GO) terms and Enzyme Commission (EC) numbers. We validate these against annotations inferred from UniProtKB/Swiss-Prot. These annotations immediately expand functional coverage for both genome and microbiome -level analyses. For organisms in FusionDB, we annotate *≥*50% more proteins than alignment to Swiss-Prot alone. For other bacteria, the gain is smaller but still reaches 30% for some organisms. For microbiomes, annotation coverage increases nearly ten-fold. Our work thus enables the analysis of microbial “transparent matter,” i.e. functions already described in earlier research that have not yet made it into usable annotations.

Our annotated Fusion function database is freely available at https://services.bromberglab.org/fusion2ai and as a reference DB for metagenome annotation https://services.bromberglab.org/mifaser.

## 1. Introduction

How much do we know about protein function? The prevailing view, shared by curators, experimentalists, and computational biologists alike, is that we know very little [1]. Experimental characterization covers a small fraction (*∼*0.2%) of known protein sequences, even when accounting for high-throughput assays [2]. Computational methods expand this space only incrementally: sequence-and structure-based approaches infer function by similarity to already-annotated molecules or their motifs/domains, rather than discovering genuinely new activities. Moreover, these methods make mistakes [3, 4, 5], propagating erroneous assignments across protein space and remaining stubbornly elusive throughout database timelines. Community assessments, such as the Critical Assessment of Function Annotation (CAFA), confirm substantial room for improvement [6]. On the other hand, literature that concerns itself with disease causing and/or phenotype-relevant genes consistently reports implicit, if not explicit, confirmations of suspected molecular mechanisms via findings of “fitting” functional annotations fished out from the sea of available data. This observation begs a question: perhaps we know more than we think and the knowledge is simply **transparent** to further research, i.e. scattered and difficult to use at scale?

We previously built the Fusion database (fusionDB), which clusters 30.6 million proteins extracted from 8,906 complete bacterial genomes into 433,891 functional groups based on function-relevant sequence similarity (HFSP; [7, 8]). This database enables reference-free organismal comparisons: organisms A and B can be quantified as 50% functionally similar even without knowing what functions they share. This design was deliberate, targeting all of bacterial functionality, with or without well-defined annotations. Indeed, we report here that of all fusionDB clusters, termed *fusion functions* from here on, two thirds (*∼*284,025 of 434K), i.e. the majority of known bacterial functional diversity, do not have even one protein with a human-readable description. Thus, only just over a third (35%; 149,866 clusters) contain any proteins with informative functional annotations, i.e. text beyond vague terms such as “hypothetical protein” or “transmembrane protein” (uncertain terms are reported in Supplementary Note S2).

Note that even the fusion functions that do contain non-vaguely annotated proteins present a challenge for practical use. Individual functions may contain distinct protein names that vary widely – a likely consequence of different source databases, annotation pipelines, and levels of curation [3, 9]. For example, function F-24979 contains members named *“dehydrogenase,” “pyrroloquinoline quinone biosynthesis protein PqqC,” “TENA/THI-4 domain-containing protein,” “CADD family folate metabolism protein,”* and *“chlamydia protein associating with death domains.”* The summary function for this cluster may be *“This protein cluster is involved in pyrroloquinoline quinone (PQQ) biosynthesis, likely playing a role in cofactor synthesis and enzyme modification in bacterial metabolic processes”*. The generation of such a description is not simply transfer of existing annotations, but an *inferred consensus function* that may not explicitly match any single member’s name. In other words, the functional identity of each cluster is latent in member gene/protein names and needs to be actively compiled.

This work aims to close this gap in **transparent** function annotations. We use a large language model (LLM; Claude [10]) to synthesize the available diverse, partial annotations of individual fusionDB functional cluster members into a single coherent function description per cluster. The key insight from our work is that while any one protein name may be incomplete or imprecise, the *consensus* across dozens to thousands of names within a functionally coherent cluster is informative. For the *∼*150K fusionDB functions that contain at least one annotated protein, we now provide an English-language description: *∼*99,000 via LLM synthesis of multiple names/annotations, and *∼*51,000 where only a single unambiguous name was available.

To bridge text descriptions with standardized vocabularies, we further used LLMs to extract Gene Ontology (GO) terms [11] from function descriptions and mapped these to Enzyme Commission (EC) numbers [12]. We further validated GO assignments against an independent reference, i.e. GO terms transferred from UniProtKB/Swiss-Prot [2] via sequence homology (HFSP*≥*14; [8]). These extracted ontology annotations make fusionDB more “quantitatively accessible” and useful for its downstream applications where comparing free text annotations on large scale is more complicated than comparing labeled abundance profiles.

Previously, we used the fusionDB functional landscape as reference for the mapping of newly sequenced bacterial genomes [13, 14] and, in mi-faser [15, 16], for annotating metagenomic reads. For both methods, users could see that a specific fusionDB function *was present* but not *what this function was*. With our new descriptions, fusionDB and mi-faser services can now return interpretable functional profiles for a large subset of common functions: roughly 84% of the functions per genome for bacterial species present in fusionDB, 47% per genome for organisms who have no relatives (as high up as class level) in fusionDB, and 26% per microbiome. For comparison, mi-faser using manually curated function annotations annotates *∼*3% of microbiome reads.

At the same time, our results clearly highlight what remains really unknown: the 284K Fusion-unannotated clusters represent only about a tenth of all proteins in our set. These are **not “transparent”** , containing no annotated members and, likely, representing the diversity of bacterial functionality that is adaptation-specific, i.e. allowing individual species or strains to adapt to the changing environment. These are smaller functional clusters with about ten proteins each, so it is no surprise that they have not yet been annotated, whether with the older homology-based models or with advanced machine learning [17, 18, 19, 20]. These proteins constitute the part of the functional “dark” matter of the bacterial protein universe that is *hidden in plain sight*. Unlike the metagenome-extracted translated ORFs [21], these proteins are more common and readily accessible, i.e. they are derived from fully assembled bacterial genomes and have known homologs. However, they still remain unannotated. Whether the current functional ontologies contain terms adequate to describe these proteins, or whether entirely new functional categories await discovery, is an open question with significant implications for our understanding of bacterial biology.

## 2. Materials and Methods

We used a combination of text-filtering, large language model (LLM) prompting, and alignment to UniProtKB/Swiss-Prot to summarize the functions of Fusion clusters in the form of (i) text-based descriptions, (ii) Gene Ontology (GO) terms, and (iii) Enzyme Commission (EC) numbers.

### 2.1. Fusion reference datasets

Fusion clusters of functionally similar proteins are derived from the complete proteomes of 8,906 distinct bacteria [7]. Here, we refer to following subsets of these proteins:

- *Full protein set* : the complete set of 30,614,981 proteins contained within any Fusion cluster. Note that the full fusionDB set of bacterial proteins had *∼*31.5M sequences, including *∼*1M singletons (no homologs in the entire set) and *∼*4K peptides shorter than 23 amino acids; these proteins were not considered here.
- *NR100* : 14,817,270 sequence-unique (at 100% identity) proteins, used to define Fusion clusters [7]
- *NR60* : the set of 3,675,307 Fusion cluster representatives (clustered at 60% sequence identity). To produce these, for each Fusion cluster individually, we ran MMseqs2 [22] *easy-cluster* with -s 7.5, –min-seq-id 0.60, –alignment-mode 3, –max-seqs 10000, –cluster-steps 2, and –cluster-reassign 1.

### 2.2. Mapping query sequences to Fusion functions

For all sets of query sequences in this work, to map proteins to Fusion clusters we ran MMseqs2 [22] with –alignment-mode 3, –num-iterations 3, -e 1e-3, –e-profile 1e-10, and -s 5.7 against either the *NR100* or *NR60* Fusion reference sets as specified in context. We then computed HFSP [8] scores, selecting as the query sequence’s Fusion assignment the one cluster with the highest HFSP*≥*14 match. At this HFSP threshold, we expect a false discovery rate of *≤*10% [8].

### 2.3. UniProtKB/Swiss-Prot alignment for EC-number and GO-term mapping

To assign EC numbers and GO terms to Fusion functions, we aligned *NR100* sequences to the Swiss-Prot database [2] (15 October 2025 release; 573,661 entries, 336,823 bacterial) using HFSP scoring and retaining the Swiss-Prot match with the highest HFSP*≥*14 for each Fusion protein. We then queried the UniProt REST API (https://rest.uniprot.org/idmapping/run; accessed December 2, 2025) to collect the GO terms and EC annotations from the *go_id* and *ec* fields from the mapped Swiss-Prot entries.

At our threshold, 5,740,960 (39%) of *NR100* sequences had a Swiss-Prot match, mapping to 264,004 unique Swiss-Prot entries. Mappings were consistent, with only 5,087 (2%) of the Swiss-Prot entries mapped to more than one Fusion cluster. However, only 17,241 of the Fusion clusters (4% of 433,891) had a Swiss-Prot match, i.e. only a small subset of Fusion functions could be annotated via homology to curated Swiss-Prot entries.

### 2.4. Data for model selection and validation

The CAFA 6 Protein Function Prediction challenge [23] tasked participants with predicting GO terms and textual function descriptions for a set of proteins. The training set for the challenge included GO term annotations from the June 18, 2025 UniProtKB release; these were experimentally validated, from a traceable author statement (evidence code TAS) or inferred by the curator (IC).

We mapped these training sequences to Fusion functions by aligning them to *NR100*. For model evaluations described here, we selected from this CAFA training set a subset of 1,000 proteins, mapping to Fusion functions with assigned Swiss-Prot entries; we limited the selection to one CAFA sequence per Fusion function. We refer to these 1,000 selected sequences as our *model-selection dataset*.

For each of the CAFA sequences, we also aligned them directly to Swiss-Prot to extract all non-self-hit matches with HFSP*≥*14; 795 of 1,000 sequences had a Swiss-Prot match.

### 2.5. Compiling function descriptions

#### 2.5.1. Preprocessing and filtering

In the original Fusion work [7], the *full protein set* was assigned to 433,891 Fusion functional clusters, with 2 to 259,895 sequences per cluster. We used the gene/protein names of these sequences, pulled from GenBank (NCBI public ftp; 28 February 2018) [24], for all work reported here. We processed these names using methods in Supplementary Material S1.

After all processing, 149,866 (35% of the 434K) Fusion functions had at least one unique *highly meaningful* name, including 32,474 (7%) that had more than five. Note that only 6% (10,064) of *∼*160K Fusion functions with at least one *certain* name, did not have any *highly meaningful* names.

We then compiled the list of each Fusion function’s protein names for LLM summarization as follows:

1. Choose the most frequent original, i.e. unprocessed, protein name for each processed *certain* name.
2. Replace any pipe (“|”) characters in these names with spaces.
3. Concatenate names in descending order by frequency of the processed name within the Fusion cluster, separating names by pipe characters.

We only prompted the LLM to produce names for the 99,093 (23% of the 434K) Fusion functions with *≥*1 unique processed name and *≥*1 *highly meaningful* name. For the 50,773 (12%) of Fusion functions containing only one unique *processed name*, which was also *highly meaningful*, we kept the most frequent unprocessed protein name as the function annotation. The 65% Fusion functions comprising proteins with only (1) *minimally meaningful* processed names or (2) uncertain names remained functionally unannotated.

#### 2.5.2. LLM function summarization

We compared function summaries generated by Haiku 3.5 (*claude-3-5-haiku-20241022* ), Sonnet 4.5 (*claude-sonnet-4-5-20250929* ), and Opus 4.6 (*claude-opus-4-6* ), all from Claude [10], for the 1,000 functions in the *model-selection dataset*. We queried all three models with identical prompts and parameters (Supplementary Note S5) using Claude’s Batch API with one request per Fusion function and each request processed independently [25]. We tuned the function summarization prompt (1) to encourage focus on the shared functionality indicated by the protein name set but, also, (2) to list unrelated functions separately without merging or inventing functional connections. The prompt gives explicit instructions to use confidence qualifiers such as ’likely,’ ’putative,’ and ’predicted’ for ambiguous cases or to state “functional details of this cluster are uncertain” when applicable. For each function, the corresponding pipe-separated list of protein names was substituted into the user prompt. No post-processing was done on the LLM-generated function summaries.

Note that Claude sometimes refused to generate responses (potential policy violations): Opus refused to generate summaries for nine functions of the *model-selection dataset* and Sonnet refused for two of the 1,000. Refused functions were kept for evaluation, contributing zeroes to most metrics.

#### 2.5.3. LLM-based GO term extraction from function descriptions

We extracted GO terms from text descriptions of protein functions using Haiku 4.5 (*claude-haiku-4-5-20251001* ), Sonnet 4.5 (*claude-sonnet-4-5-20250929* ), and Opus 4.6 (*claude-opus-4-6* ) to both validate the function summaries and, potentially, provide GO terms for the *∼*136K (91% of *∼*150K) Fusion functions with textual function descriptions but no GO annotations extractable from Swiss-Prot matches.

For this task, we used a system prompt in addition to a user prompt (Supplementary Note S7) to give the model a role (“a cautious expert bioinformatics curator”), provide a mix of output examples, and clearly specify the output format. None of the examples we used were from functions in the *model-selection dataset*. Though we requested the model to assign low, medium, and high reliability to each extracted GO term, we used the union across all three reliability levels unless otherwise specified (see Table S7 for a comparison across reliability levels).

We post-processed model outputs to replace obsolete IDs with their current equivalents from the June 1, 2025 release of the *go-basic* graph and discarded any IDs that were neither primary nor known alternate terms in that release (i.e. likely hallucinated).

### 2.6. Evaluating function descriptions

#### 2.6.1. LLM as a judge

We evaluated the different Claude models’ function summary abilities using LLM-as-a-judge. That is, we asked the Haiku 4.5 judge for the likelihood that a given, specific protein function is captured by the text describing its Fusion function. We prompted *claude-haiku-4-5-20251001* to score (range 0 to 10) ten unprocessed protein names for each of the 1,000 functions of the *model-selection* set (labeled as *certain*, randomly sampled, with replacement; prompt in Supplementary Note S6). As an evaluation control, for each protein name we also prompted the model with a functional summary of another, randomly selected Fusion function (*wrong-function summaries*). Note that we expected Haiku 4.5 to be sufficient as a judge because although the task required a strong scientific vocabulary, it is a relatively simple measure of semantic similarity. Haiku 4.5 judge provided scores for each query, except for fewer than ten cases per summarization LLM; for these we set their likelihood =0.

We performed paired, two-sided t-tests comparing likelihoods that protein names were correctly assigned to their function summaries vs. wrong-function summaries. The judge-estimated likelihood differences were significant (*p <* 0.05) across most function summaries generated by the LLMs (975 of 1,000 Haiku 3.5, 984 of 998 Sonnet 4.5, and 984 of 991 Opus 4.6). The 95% confidence interval for the mean *difference* in likelihood of correct vs. wrong function summaries, was 6.44-6.54 for Haiku 3.5, 6.71-6.81 for Sonnet 4.5, and 6.83-6.93 for Opus 4.6. We additionally measured each model’s performance via an area under the receiver-operator curve (AUROC; per query, positives are matches to correct function summary and negatives are matches to a wrong summary) for LLM-as-judge score thresholds from 0.5 to 9.5 in steps of 1.0, attaining 0.86 for Haiku 3.5, 0.80 for Sonnet 4.5, and 0.79 for Opus 4.6.

While the mean likelihood difference for correct vs. wrong function summaries increased with model complexity as expected, AUROC was slightly better for Haiku, i.e. the simplest model. Two factors likely contribute to this observation. First, model prompt refusals (nine for Opus 4.6, two for Sonnet 4.5, none for Haiku 3.5) contributed zeroes that lowered only the more complex models’ AUROC scores. Additionally, the more capable models likely produced richer descriptions that could occasionally also fit a wrong cluster’s proteins, increasing tail overlap that AUROC penalizes. Given the small magnitude of these differences, however, all models can be said to perform similarly. As such, we used the cheaper Haiku 3.5 functional summaries of Fusion functions.

#### 2.6.2. Evaluating LLMs for functional text processing

We used the CAFA ground truth GO term designations to evaluate (Supplementary Materials S8) different LLMs’ abilities to extract GO terms from text. For each protein (**P**) in the *model-selection dataset*, mapped to a Fusion function (**F**), we assigned GO terms to **P** using the following homology (1-3) and LLM-based (4) methods:

1. *Fusion-SP* : Align all Swiss-Prot proteins to Fusion function representative sequences (*NR100* ), selecting for each the Swiss-Prot entry with the highest HFSP*≥*14 match. To each **F**, assign the union of GO terms from all Swiss-Prot matches. Each **P** in **F** inherits this assignment
2. *All SP Matches*: Use the union of GO terms from all Swiss-Prot proteins aligned to **P** with HFSP*≥*14, excluding self-hits
3. *Best SP Match*: Use GO terms from the one Swiss-Prot protein aligned with highest HFSP*≥*14 to **P**, excluding self-hits
4. *LLM Text, LLM GO* : Use a Claude model to extract GO terms from the function summary generated for **F**. Each **P** in **F** inherits this assignment

No approach was perfect at extracting GO terms from functional descriptions that would match the proteins’ CAFA-assigned gold standard GO terms (Tables S4, S5, S6). For LLM-based designations, as perhaps expected, the choice of Claude model for GO term extraction had a larger effect on performance than the choice of model for function summary generation. This was evidenced by the substantial improvement in performance when using Opus-4.6 over Haiku-4.5 for extraction of GO terms from Haiku-3.5-generated summaries (Wang similarity, Eq. 2, = 0.52 [95% CI: 0.51-0.54] vs. 0.45 [0.43-0.47]), but minimal improvement for using Opus-4.6 vs. Haiku-3.5 for summary generation (with Opus-4.6 GO term extraction; Wang similarity = 0.55 [0.53-0.57] vs. 0.52 [0.51-0.54]). Our performance evaluation results thus confirm the above observation that Haiku 3.5 is sufficient for function summarization. Note that it was further preferred for our analysis due to its lower cost (and likely energy consumption) of $44 for extracting 99,093 function summaries versus $164 for Sonnet 4.5 and $274 for Opus 4.6. Note that over the duration of our research period, Anthropic released Opus 5, which exhibited slightly better performance than Opus 4.6 for GO term extraction on our *model-selection dataset*.

#### 2.6.3. Validating LLM-extracted Fusion functional summaries via GO term mapping

We compared the LLM (Opus 5) ability to extract GO terms (Supplementary Note S7) from a given protein’s (1) Swiss-Prot protein name(s), (2) gold standard, curator-generated function descriptions, i.e. CC id in Swiss-Prot entry, and (3) corresponding Haiku 3.5-generated Fusion function description. Here, we considered the set of 795 Swiss-Prot entries that were best sequence matches (with HFSP*≥*14) for proteins in the *model-selection dataset*, self-hits excluded. We queried the UniProt REST API (March 11, 2026) for their “protein name(s)” (*DE RecName:*) and “function” (*CC -!- FUNCTION:*) fields. All 795 had names, but only 647 had function descriptions. Fusion functions (and corresponding functional summaries) were assigned to these proteins at the compilation of the *model-selection set*. All three sets of LLM-extracted GO terms corresponding to the different function description types were matched to the ground truth GO terms mined from the corresponding Swiss-Prot entries (*DR GO;*).

We evaluated GO term mapping performance across 1K bootstrap iterations sub-sampled at 60% without replacement from this protein set (Table S1; full per-LLM model results are in Table S2). GO metrics were better for name/curated text -based function descriptions vs. LLM-based ones for the MF subontology, although the differences from curated function text were low. Other metrics and evaluation of other subontologies (BP and CC) deemed free text descriptions comparably informative of the corresponding GO terms. In other words, GO terms extracted from the LLM functional summaries were comparable to those mapped from manually curated Swiss-Prot *protein names* and *function descriptions*.

To assess whether LLM-judge confidence predicts annotation accuracy, we scored each functional summary as described above (2.6.1). That is, for every protein in a given function, we asked the judge to rate — from the protein name alone — how likely it was that the function summary described that protein. Averaging these ratings over the 10 proteins per function gave a single confidence score per function summary. We then correlated that score with the accuracy of the GO terms extracted from the same summary. GO-term extraction accuracy was measured as Wang and Jaccard (Supplementary Note S8) similarity between the extracted GO terms and corresponding Swiss-Prot annotations, computed separately for the BP, MF, and CC GO subontologies. Correlations were evaluated with two-tailed Spearman and Pearson tests (Table S3). LLM’s functional summarization confidence correlated with GO term extraction accuracy weakly if at all; correlation was significant only for the BP subontology.

We note that the functional diversity within a Fusion function is the upper bound for the accuracy that any method can reach when extracting GO-term annotations, since the LLM cannot match the ground truth more closely than the function’s own proteins match each other. To evaluate LLM extraction of GO terms against this empirical ceiling and a cross-function baseline, we computed, for each Fusion function, three pairs of quantities: (1) the mean and worst pairwise GO-term similarity across all pairs of Swiss-Prot proteins mapped to this function (the ceiling), (2) the mean and worst similarity between the function’s LLM-extracted GO terms and those same Swiss-Prot proteins’ GO terms (our method), and (3) the mean and worst similarity between the function’s LLM-extracted GO terms and the Swiss-Prot proteins’ GO terms of a randomly sampled other Fusion function (the cross-function baseline). We used only experimentally validated GO terms (evidence codes EXP, IDA, IPI, IMP, IGI, and IEP, and their high-throughput equivalents), retrieved from the UniProt REST API (https://rest.uniprot.org/uniprotkb/search; accessed April 21, 2026). We compared the Wang similarity distributions of our method against the ceiling with two-sample Kolmogorov–Smirnov (KS) tests and expected the MF terms to be best outlined because Fusion functions are aimed at capturing molecular functional similarity [7]. While our method was statistically distinguishable from the ceiling (MF: *D* = 0.138, *p* = 6.43*E −* 8; BP: *D* = 0.150, *p* = 6.7*E −* 9; CC: *D* = 0.357, *p* = 2.9*E −* 9), it was much closer to the ceiling than to the baseline, especially for the MF and BP subontologies (Figure 1; Figure S1).

**Figure 1:**
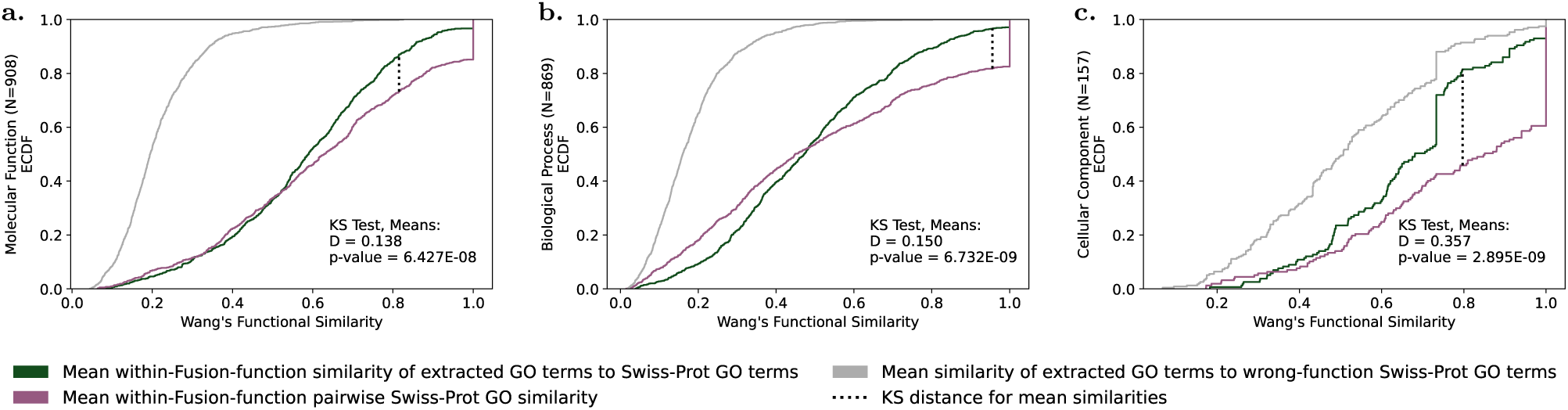
Variety of GO terms within a Fusion function bounds maximum accuracy of text-derived GO terms. Empirical cumulative distribution functions (ECDFs) of per Fusion function (1) within-function pairwise similarities of experimentally validated Swiss-Prot GO terms (purple) and (2) similarities between GO terms extracted by Opus 5 from Haiku-3.5-generated function summaries and Swiss-Prot GO terms for the corresponding Fusion function (dark green) and a randomly sampled other Fusion function (gray) are shown, and lower lines indicate higher accuracy. All Fusion functions with GO terms from at least two Swiss-Prot entries and at least one GO term extracted with Opus 5 from the Haiku-3.5-generated function summary per subontology are considered. Kolmogorov-Smirnov tests (at p-value cutoff of 0.05) indicate that the within function GO term variety vs. text-derived GO term variety distributions are significantly different, though they are noticeably closer to each other than the wrong-function baseline especially for MF (a) and BP (b) GO terms.

### 2.7. Data sets for coverage analysis

We measured the coverage of proteomes/genomes/metagenomes with Fusion annotations for the following samples (UniProt proteome or MGnify genome/proteome IDs in parentheses): *M. mycoides* (UP000682489; unpublished), *Ca. N. bennettiae* (UP000248636; [26]), *Ca. M. zealandia* (UP000514400; [27]), *D. mccartyi* (UP000053577; [28]), *Ca. Poseidoniales* archaeon (UP000724444; [29]), and a Binatia metagenome-assembled-genome (MGYG000523568; [30]). Additionally, we annotated Asgard archaea (kingdom Promethearchaeati) proteins translated from metagenome-assembled-genomes (NCBI BioProject PRJNA1050611; [31]). While assemblies from two of the above species are in the FusionDB reference dataset, none of the genome assemblies we evaluate here were themselves in FusionDB. To estimate previously available annotation coverage, we aligned thus-collected protein sets to Swiss-Prot with MMseqs2 (same parameters as above; 2.2) to evaluate how many had Swiss-Prot homologs with E-value*<* 10*^−^*^3^ and with the stricter functional similarity requirement of HFSP*≥* 14.

We also computed the fractions of reads from annotated Fusions across six metagenome samples from the Deepwater Horizon oil project, including pre-oil (SRR1566021, SRR1569462), oil (SRR1569742, SRR1569812), and post-oil recovery (SRR1570801, SRR1570802) samples (NCBI BioProject PRJNA260285; [32]). We compared Fusion annotation coverage vs. the fraction of reads that *mifaser* could assign E.C. numbers for, using the *mifaser* web application with the GS-24-all reference database and quality control on (accessed August 26, 2026). Quality control was done with *fastp* v0.20.1 [33] with *phredquality* = 20 and *readlength* = 40.

### 2.8. Web application

The Haiku-3.5-generated text summaries for Fusion clusters, along with EC numbers and GO terms retrieved from Swiss-Prot, are available through the *mifaser* web application (https://services.bromberglab.org/mifaser [15]) via the *fusion-v2* reference database option. The *mifaser* web application translates and aligns query reads against the *NR60* Fusion reference set and provides a user-friendly table for searching and filtering through the annotated Fusion functions. Exploration and comparison of Fusion profiles across organisms with both GO terms from Swiss-Prot and LLM-extraction (Opus 5) is available through the *Fusion2AI* web application (https://services.bromberglab.org/fusion2ai).

## 3. Results and Discussion

### 3.1. Annotation landscape of Fusion functions

Of the 433,891 Fusion functions, only 149,866 (35%) contained at least one protein with a *highly meaningful* name **(Methods)** and could thus be assigned a text description (Table 1). These include 99,093 (23%) functions summarized by the LLM from multiple names and 50,773 (12%) annotated with a single unambiguous name. The remaining 284,025 functions (65%) were left without descriptions: 10,334 (2%) carried only *minimally meaningful* names and 273,691 (63%) had no *certain* names at all. Of 810,817 Fusion singletons, 53,100 (7%) had a *highly meaningful* name; note that we do not search singletons by default in fusionDB settings. As expected [20], larger functional clusters tended to be annotated, whereas functions without *highly meaningful* names were, on average, smaller (Figure 2). This observation has at least two explanations: (1) technically, larger protein groups are simply more likely to have at least one annotated protein and (2) scientific interest skew assures that sparsely populated, adaptation-specific groups consistently escape annotation. We note, however, that “true functional discovery” is likely to lie in these poorly explored spaces.

**Figure 2:**
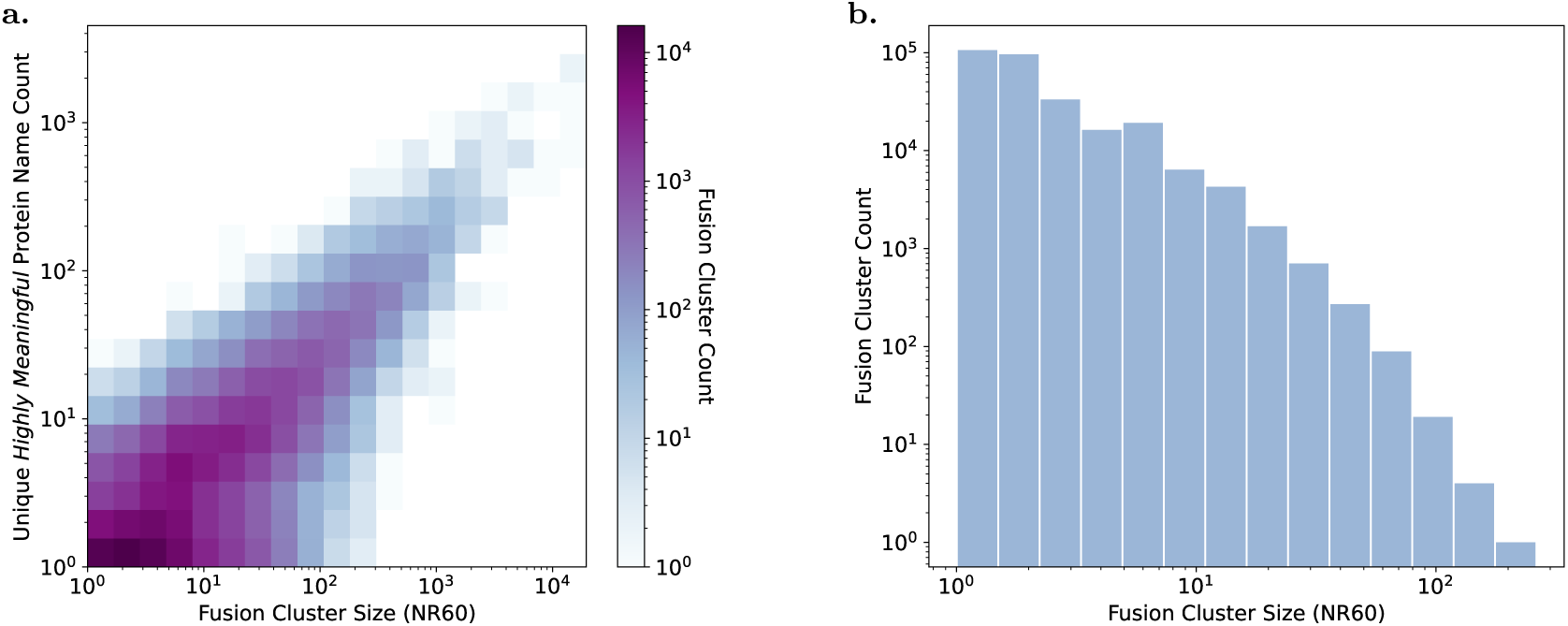
The number of *highly meaningful* protein names in a Fusion cluster’s reference set generally increases with cluster size (a), and clusters without *highly meaningful* names are smaller on average (b). Cluster size here is the count of reference members in the non-redundant set clustered at at 60% sequence identity (*NR60* ). (a) covers the 149,866 Fusion clusters with any *highly meaningful* names, and (b) covers the 284,025 without any *highly meaningful* names.

**Table 1:** Counts of Fusion functions with different types of annotations.

| Text Annotation | Fusion Functions | Swiss-Prot GO* |  |
| --- | --- | --- | --- |
|  |  | Count |  |
| LLM-Summarized | 99,093 | With | 13,315 |
|  |  | Without | 85,778 |
| Single *Highly Meaningful* Name | 50,773 | With | 715 |
|  |  | Without | 50,058 |
| Only *Minimally Meaningful* Names | 10,334 | With | 113 |
|  |  | Without | 10,221 |
| No Certain Names | 273,691 | With | 445 |
|  |  | Without | 273,246 |
| Total | 433,891 | With | 14,588 |
|  |  | Without | 419,303 |

\* With: Functions that have at least one *NR100* protein that matches (with highest HFSP $\geq 14$ ) a Swiss-Prot entry with GO annotations at any evidence level.
We labeled names of proteins in each Fusion cluster’s reference set as *certain* or *uncertain* and *highly meaningful* or *minimally meaningful* (Section 2.5.1). Only $\sim 150\text{K}$ functions (35%) had any *highly meaningful* names, and about a tenth of that amount (3.4%) had Swiss-Prot homologs with GO annotations.

Similarity to functionally labeled proteins in Swiss-Prot, a large repository of available curated annotations, covered only a small fraction of the complete FusionDB landscape: just 14,588 (3%) of all Fusion functions had any GO annotations transferable by homology. These were concentrated within the LLM-summarized functions (13,315 of 14,588; Table 1). Thus, *∼*136K (91% of the *∼*150K) text-described functions had no Swiss-Prot-derived GO terms, defining the gap that our text-to-GO extraction aimed to fill.

### 3.2. LLM generates coherent and correct function descriptions

Function descriptions generated through our pipeline are specific, precise, and often easier for a broad audience to understand than long lists of protein names often containing abbreviations, e.g. “VapB4”, “LppE”, “YVMC”, etc. We showed that LLM-summarized function descriptions are specific to the proteins they describe rather than overly general. Furthermore, GO terms extracted from functional summary texts were nearly as precise (*∼*0.8 vs. *∼*0.85; Supplementary Note S8) as those extracted from Swiss-Prot high-quality, curated text of protein names or function descriptions (Table S1).

Often, the GO terms extracted from a text summary were ancestors of the GO terms carried by the cluster members proteins’ Swiss-Prot entries. We took this to mean that our functional descriptions reasonably captured the shared functionality per Fusion cluster. Recall that Fusion clustering is based on a sequence similarity and cutoffs chosen to reflect functional similarity (HFSP; [8]). Thus, even sequence-diverse proteins within a cluster are likely to share the same high-level molecular function or to occupy similar roles in a common biological process.

For example, F-9 is a large cluster with *∼*4K (*NR60* ) members, corresponding to 507 unique Swiss-Prot proteins. Its LLM-summarized function description is: “[t]he protein cluster primarily functions in aldehyde oxidation and dehydrogenase activity, with NAD(P)+ as a cofactor, catalyzing oxidative decarboxylation and metabolic transformations of various aldehydes and semialdehyde compounds across bacterial metabolic pathways.” From this description we extracted the molecular function (MF) GO term “oxidoreductase activity, acting on the aldehyde or oxo group of donors, NAD or NADP as acceptor” (GO:0016620; high reliability).

Of the 90 (18% of 507) Swiss-Prot proteins that have MF GO terms and map to F-9, six match (the high-reliability term) GO:0016620 exactly and a further 77 proteins carry 33 of GO:0016620’s unique descendants (through “is_a” relationships). The other seven Swiss-Prot F-9 hits carry GO terms describing other oxidoreductase activities or only binding preferences. Of the proteins with GO:0016620 descendant annotations, only four are assigned additional other functions. GO:0016620 thus strikes a reasonable balance between specificity and generality for describing the cluster’s function. The proteins’ binding targets, by contrast, were highly diverse – spanning hormones, nucleic acids, lipids, proteins, and small molecules. Therefore, binding-related functionality was not mentioned in the summary, nor extracted as GO terms. This pattern explains the lower evaluated performance for molecular function than for biological process (MF < BP) in GO term extraction from function text. It is also consistent with evolutionary conservation of biological processes even under environment-driven molecular function adaptation.

We aimed to reduce LLM hallucination in functional descriptions by labeling protein names as *highly* vs. *minimally meaningful* and only using LLM summarization when there was at least one *highly meaningful* name. Our prompt also clearly stated to “NEVER invent functions for unrecognized proteins.” However, we still observed some cases where function descriptions were not directly supported by the input list of protein names.

For example, F-58152 has only two unique, *certain* protein names: “zinc-binding protein” and “transmembrane zinc-binding protein.” The generated function description is “[t]hese zinc-binding proteins likely facilitate zinc transport and metal homeostasis across bacterial cell membranes, potentially regulating cellular zinc uptake and distribution.” While it is quite possible that the functional cluster could be involved in zinc transport, as suggested by the function description, it could also be involved in signal transduction, such as with a chemoreceptor zinc-binding (CZB) domain [34]. The LLM similarly made a logical leap for F-144702, which has just the names “conserved repeat domain protein” and “parallel beta-helix repeat protein” but summarized function description: “[t]he proteins likely participate in structural stabilization and protein-protein interactions through conserved repeat domains, potentially involved in bacterial cellular processes.” “[P]arallel beta-helix repeat protein” would have been more appropriately labeled a *minimally meaningful* name, in which case these names would not have been summarized by the LLM through our pipeline, but the word “parallel” is not in the *low-content vocabulary* (Supplementary Note S4) because it has meanings that may not be structural (e.g. “in parallel with”).

Such hallucinations were not pervasive. In our LLM-as-a-judge evaluation (2.6.1), across all functions (100%), the mean likelihood of a protein name falling under the wrong function summary was *≤*4/10, while for correct function summaries only 23 (2.3%) of mean likelihoods in the same range. This observation supports our conclusion that functional summaries were not overly broad or often erroneous. We also note that only about a tenth (*∼*13%) of MF GO term sets extracted from function summaries were dissimilar to Swiss-Prot annotations (2.6.3, Figure S1; Wang similarity *≤*0.3, a level at which, generally, only higher level GO terms are shared). Note that sparsity of Swiss-Prot annotations and outdated GO training data, as much as hallucination during GO extraction, could have contributed to these cases (Supplementary Data; condensed lists of raw protein names per cluster and LLM function summaries).

We also note that the LLM did, in some cases, call out unclear or conflicting annotations. For example, F-58749 had protein names “BRCA1,” “group-specific protein,” and “BRCA1-like protein.” The function description was appropriately: “[f]unctional details of this cluster are uncertain. These protein names suggest potential mammalian/human-associated proteins, not a bacterial protein cluster, which requires further clarification.”

### 3.3. GO terms and EC numbers extracted from text descriptions

LLM-based GO term extraction (Haiku 3.5 summaries, Opus 5 GO term extraction; **Methods**) annotated 134,060 Fusion functions (89% of all text-annotated *∼*150K), the majority of which (*∼*136K) previously had no GO annotations or even sufficiently close annotated homologs (in Swiss-Prot). Extracted terms spanned all three GO subontologies: 100,585 functions had MF GO terms, 109,225 had BP GO terms, and 30,717 had CC GO terms. Opus 5 refused to extract GO terms for 187 functions due to potential policy violations and reached its token limit for 213 functions. Each batch of 10K requests for GO extraction took 16-36 min. and cost an average of $9 (USD) with ephemeral (5-min.) caching of the system prompt (Table S8).

We further mapped extracted molecular function GO terms to Enzyme Commission (EC) numbers [35], yielding 206 unique third-level EC assignments for 53,658 functions. As expected, EC coverage was lower than GO coverage, since only catalytic clusters receive EC numbers. These text-derived GO and EC annotations, together with those transferred from Swiss-Prot, constitute the machine-readable layer of the annotated Fusion database.

### 3.4. Validation against Swiss-Prot homologs

LLM function summary confidence was only weakly associated with the similarity of extracted GO terms vs. Swiss-Prot homolog curated ones. For biological process terms the correlation was significant but weak (Spearman’s *ρ* = 0.22, *p* = 5 *×* 10*^−^*^10^; Pearson’s *r* = 0.18, *p* = 3.6 *×* 10*^−^*^7^). It was weaker for molecular function terms (*ρ* = 0.04, *p* = 0.26; *r* = 0.08, *p* = 0.02) and not significant at all for cellular component terms. These observations suggest that LLM confidence is not a useful proxy for annotation accuracy. The decoupling may be due to the many alternative phrasings that can be used to describe a single GO term (text understanding failure), as well stem from the model’s difficulty in capturing functions that are implied rather than stated explicitly by protein names.

Furthermore, in pairwise comparisons of within-function MF GO term annotations, we found that Swiss-Prot annotated GO term similarities vs. the LLM-GO to Swiss-Prot-GO similarities were close with a maximum difference in their ECDFs of *D* = 0.138 around a Wang similarity of 0.8 despite being distinguishable (KS test) (Figure S1). Functional diversity within a Fusion cluster indicates the limit to the accuracy of any GO extraction method, i.e. the LLM cannot match the ground truth better than the ground truth protein annotations match each other. Indistinguishability of the distributions would indicate that LLM-extracted molecular function GO terms are about as accurate as the intrinsic, experimentally observed diversity allows. Note that the experimentally observed diversity does not necessarily reflect the true functional diversity within a function cluster, since annotations are sparse and the annotated proteins are not a random sample. However, curation tends to add proteins similar to those already annotated rather than novel ones [20]. The observed within-cluster diversity is therefore unlikely to grow substantially as databases expand, and the ceiling we measure should remain a stable reference. If anything, the unobserved diversity would lower the true ceiling, making our reported performance to it a conservative estimate.

### 3.5. Functional annotation coverage expands for fusionDB and mi-faser

Because Fusion functions underlie both whole-genome mapping (fusionDB) and metagenomic read annotation (mi-faser), our new descriptions directly increase the share of each query that can be assigned a human-readable function. For a typical bacterial genome, proteins mapped to a Fusion function previously returned only a cluster identifier. With our annotations, 93% of a genome’s Fusion-mapped proteins (measured as a mean over five bacterial genomes) now carry an English-language description, up from 70% via Swiss-Prot homology alone. For species included in our reference, more of the proteins could be annotated with Fusion text than had any Swiss-Prot homologs with E-value*<* 10*^−^*^3^. Although Archaea are not in fusionDB, we were also able to annotate somewhat more proteins across two archaeal sets than there were functionally similar proteins in Swiss-Prot (HFSP*≥*14; Figure 3).

**Figure 3:**
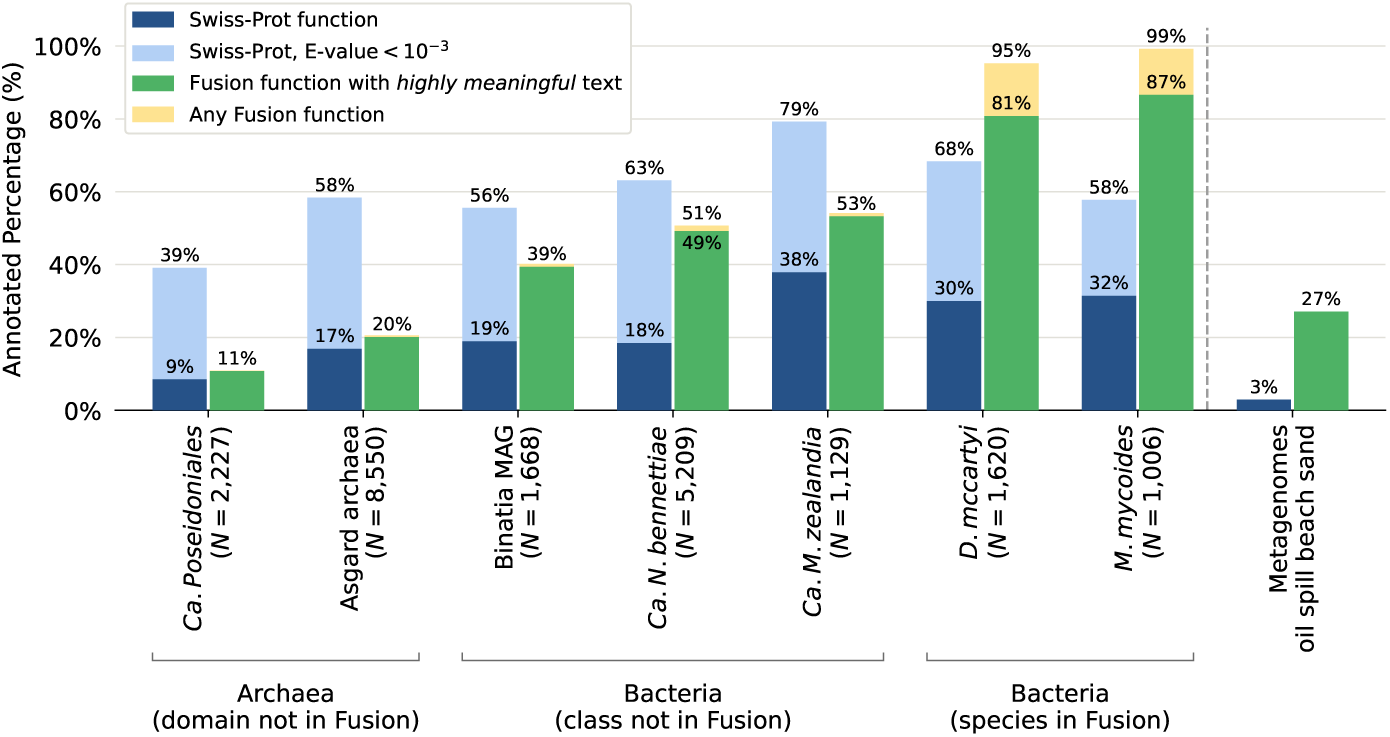
Fusion expands functional annotation at the genome and metagenome levels. More proteins are mapped to Fusion functions with *highly meaningful* text annotations (green) than are functionally matched to Swiss-Prot (dark blue; *HFSP ≥* 14 for proteomic data; mapped to mi-faser’s [15] *GS-24-all* database for metagenomes) across all samples (2.7). Each sample along the x-axis to the left of the dashed line corresponds to one proteome except for Asgard archaea, which is a protein set containing multiple metagenome-assembled-genomes. The oil spill metagenome bars represent means across six metagenome samples (Table S9). Light blue bars show any significant Swiss-Prot matches (MMseqs2 with E-value cutoff of 10*^−^*^3^). Yellow bars represent matches to any Fusion functions and do not have labeled percentages when their visible portions are *<* 1% for proteome samples. Fusion singletons are included but make up *<* 2% of matches for species in Fusion and *<* 0.5% of matches for all other samples. *N* is the the number of protein sequences in the sample.

The gain is larger for metagenomes, where reference-free clustering matters most. Across six microbiome samples, the fraction of mi-faser–annotated reads with an interpretable function rose from 3% to 26% (Figure 3; Table S9).

### 3.6. The unannotated functional dark matter

Our pipeline leaves 284,025 Fusion functions (65%) without a text description – the 273,691 (63%) with no *certain* names and the 10,334 (2%) with only *minimally meaningful* names. Unlike metagenome-derived “dark matter,” the proteins in these clusters are drawn from fully assembled bacterial genomes and have known homologs, yet remain functionally opaque. The clusters are small (*∼*10 proteins per cluster; Figure 2) and together account for only about a tenth of all proteins in our set. However, their persistence despite decades of homology-and machine-learning-based annotation suggests they represent genuinely under-characterized, likely adaptation-specific bacterial functions rather than an artifact of missing names.

This residue frames both the reach and the limits of our approach. The central lesson is that an LLM can act as an aggregator of noisy, partial annotations: while any single protein name may be incomplete, the consensus across a functionally coherent cluster is informative enough to yield a specific, validated description. That leverage, however, requires at least one *highly meaningful* name per cluster – exactly what the dark-matter clusters lack – so no amount of summarization can annotate them. Our results should be read with three limitations in mind: residual LLM hallucination (mitigated but not eliminated by our meaningfulness filtering), the variable quality and 2018 vintage of the underlying GenBank names, and the restriction of both the reference and the analysis to bacteria.

These same limitations point to clear next steps: refreshing the underlying genome and name sets to a current release, extending the framework to archaea and eukaryotes, and testing whether existing GO/EC vocabularies can even describe the dark-matter clusters or whether new functional categories are required. Whether these “hidden in plain sight” proteins encode known functions awaiting the right words or novel biology awaiting discovery remains, for now, an open and consequential question.

## Data Availability

All data are available in the main text, the Supplementary Materials, or referenced permanent online data repositories. Fusion function annotations for functions in the *model-selection dataset* or the pairwise evaluation (Section 2.6.3; Figures 1 and S1), Swiss-Prot alignment data, and predictions for the full CAFA 6 bacterial training set: 10.5281/zenodo.23022359. Code: bitbucket.org/bromberglab/fusion-v2-annotation. Fusion reference sequences and their Fusion function mappings along with the protein names used in this paper were previously published [7] and are available at 10.6084/m9.figshare.21599544 and 10.6084/m9.figshare.23826462.

## Acknowledgements

We thank R. Prabakaran and J.Liu (both Emory University) for insightful discussions on the utility of approximated (English language) protein function summaries. We are sincerely thankful to Y. Mahlich (Pacific Northwest National Labs) for helping us understand the Fusion functional clustering decisions and data repository structure. We thank Edgar Leon and Sergio Gramacho (both Emory) for help with the maintenance of Bromberglab computational resources that allowed for the development and maintenance of the Fusion2AI and specifically to Sergio Gramacho for guidance in HPC application design. We are exceedingly grateful to all scientists who contribute their work and understanding to functional databases.

## Author Contributions

**A.MB.**: Data Collection, Data Analysis, Writing - Original Draft, Writing - Review & Editing. **Y.B.**: Conceptualization, Methodology, Writing - Original Draft, Writing - Review & Editing, Funding and Resources.

## Declaration of Generative AI and AI-assisted Technologies in the Manuscript Preparation Process

During the preparation of this work the authors used (Anthropic) Claude [10] in order to format figures, write/analyze code, and draft help pages for web tools as well as Google’s Gemini for graphical abstract creation. After using these tools/services, the authors reviewed and edited the content as needed and take full responsibility for the content of the published article.

## Supplementary Data

### Note S1. Text Processing and Filtering

**First,** we used rule-based methods to label protein names as *certain* (e.g. “Translation initiation factor IF-2” or “Dihydrouridine synthase family protein”) or *uncertain* (e.g. “hypothetical protein” or “putative phosphoesterase”). We removed uncertain protein names from further processing. Fusion functions with no *certain*-named proteins were left without text descriptions.

*How?* We preprocessed the text by lowercasing and replacing all punctuation with spaces and then fuzzily matched against our *uncertainty vocabulary* (Supplementary Note S2). All protein names containing a word, phrase, or prefix in the uncertainty vocabulary or starting with ’potential’ were marked uncertain unless the words ’substrate’ or ’specificity’ occurred immediately adjacent to the vocabulary term. Here, we aimed to preserve informative protein descriptions even when the substrate or binding specificity was unknown, e.g. “ABC transporter ATP-binding protein - unknown substrate” was *certain*, but “putative ABC transporter” was *uncertain*. A word was considered a fuzzy match to the vocabulary if its Damerau–Levenshtein edit distance (calculated with the NLTK metrics.edit_distance function [36] with the transpositions parameter set to True) was at most =1 for every 6 letters in the vocabulary term. For multi-word phrases in the uncertainty vocabulary, word-by-word fuzzy matching was done. With this processing, 273,691 Fusion functions (63%) were left with no *certain* names.

**Then**, we summarized protein names remaining in each Fusion cluster using an LLM. To combat hallucinations for clusters that contained only overly broad protein names (e.g. “helix-turn-helix domain protein” or “outer membrane protein”), we used another set of rules to label names as *highly meaningful* or *minimally meaningful*. We only used LLMs to summarize functions with at least one *highly meaningful* name.

*How?* For each Fusion function we condensed its set of unique names by mapping closely related phrases to more general terms, e.g. ’superfamily’ to ’family’ and ’outer membrane’ to ’membrane’ (Supplementary Note S3), and removing the terms ’conserved,’ ’domain,’ ’domain containing,’ ’domains,’ ’family,’ and ’precursor’. If more than one protein had identical names post-processing, we retained only one per cluster.

We considered a thus-processed name to be *highly meaningful* if it contained any words that were not in our *low-content vocabulary* (Supplementary Note S4), which was manually curated from the collection of frequent uni and bi -grams in our function names. The *low-content vocabulary* included structural motif words, amino acid names and three-letter codes, text for numbers one through ten, and other manually curated low-content words. To match these words, we again used fuzzy matching (Damerau–Levenshtein edit distance to vocabulary term of *≤* 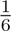 ). We retained some names with low-content matches based on domain knowledge ( “Meaningful fuzzy matches” in Supplementary Note S4). We only considered amino-acid references to be *minimally meaningful* if they were followed by the word ’rich’ with only other amino-acid names or the word ’and’ in between; e.g., we considered “alanine and glycine rich transmembrane protein” to be a *minimally meaningful* protein name.

### Note S2. Uncertainty Vocabulary

**Uncertainty words**

“candidate”, “duf”, “fragment”, “homolog”, “homologous” “homologue”, “hypothetical”, “hypothetically”, “incomplete”, “inferred”, “partial”, “possible”, “possibly”, “potentially”, “predicted”, “presumable”, “presumably”, “presumed”, “presumptive”, “presumptively”, “probable”, “probably”, “provisional”, “provisionally”, “putative”, “putatively”, “suggested”, “suggestion”, “suspected”, “tentative”, “unannotated”, “unassigned”, “uncertain”, “uncharacterised”, “uncharacterized”, “unclassified”, “unconfirmed”, “undefined”, “undetermined”, “unknown”, “unnamed”, “unverified”

**Uncertainty phrases**

“complete genome”, “high potential”, “not available”, “not classified”, “not yet annotated”

**Uncertainty prefixes**

“duf”

**Uncertainty first words**

“potential”

**Okay uncertainties (can immediately precede or follow a term indicating uncertainty; e.g., a protein with “unknown substrate” will not be counted as uncertain despite including “unknown” in its name)**

“specificity”, “substrate”

### Note S3. Term Condensing

**Condensed to “membrane”**

“inner membrane”, “innermembrane”, “integral membrane”, “membrane associated”, “membrane bound”, “membrane flanked”, “membrane spanning”, “outer membrane”, “outermembrane”, “transmembrane”

**Condensed to “family”**

“sub family”, “subfamily”, “super family”, “superfamily”

### Note S4. Low-Content Vocabulary

**Stopwords and stopphrases**

“conserved”, “domain”, “domain containing”, “domains”, “family”, “precursor”

**Low-content words**

“an”, “anchor”, “anchored”, “and”, “associated”, “bacterial”, “bacteriophage”, “basic”, “bifunctional”, “binding”, “by”, “c”, “capsule”, “cell”, “cellular”, “class”, “cluster”, “complex”, “complexity”, “component”, “components”, “containing”, “contains”, “cytoplasmic”, “cytosolic”, “dual”, “element”, “elements”, “enzyme”, “export”, “exported”, “expressed”, “external”, “factor”, “for”, “from”, “function”, “gene”, “genes”, “general”, “group”, “head”, “hydrophobic”, “in”, “includes”, “including”, “integral”, “intracellular”, “involved”, “kda”, “large”, “layer”, “ligand”, “like”, “long”, “low”, “major”, “member”, “membrane”, “meta”, “minor”, “miscellaneous”, “motif”, “multi”, “multicomponent”, “multipass”, “multiple”, “n”, “non”, “not”, “of”, “on”, “or”, “origin”, “ortholog”, “other”, “paralog”, “part”, “periplasmic”, “peptide”, “phage”, “preprotein”, “product”, “prophage”, “protein”, “proteins”, “receptor”, “region”, “related”, “repeat”, “repeats”, “repetitive”, “rich”, “secreted”, “sensor”, “sequence”, “sh3”, “short”, “signal”, “small”, “spanning”, “specific”, “subunit”, “subunits”, “surface”, “tail”, “target”, “terminal”, “the”, “to”, “truncated”, “type”, “wall”, “with”

**Number text**

“one”, “two”, “three”, “four”, “five”, “six”, “seven”, “eight”, “nine”, “ten”

**Structural motif words**

“aba”, “aligned”, “alpha”, “alphabeta”, “arm”, “armadillo”, “bab”, “barrel”, “bba”, “beta”, “bhlh”, “bridge”, “bundle”, “cage”, “chain”, “clam”, “coil”, “coiled”, “complex”, “dimer”, “distorted”, “double”, “down”, “fibrous”, “fold”, “globular”, “greek”, “hairpin”, “helical”, “helix”, “hhh”, “hlh”, “hook”, “horseshoe”, “hotdog”, “hth”, “jelly”, “jellyroll”, “key”, “knot”, “layer”, “loop”, “modular”, “monomer”, “multimer”, “nest”, “niche”, “non”, “oligomer”, “orthogonal”, “pattern”, “plug”, “prism”, “propeller”, “ribbon”, “roll”, “sandwich”, “sheet”, “single”, “solenoid”, “super”, “tetramer”, “trefoil”, “trimer”, “triple”, “turn”, “type”, “up”

**Amino acids**

“alanine”, “arginine”, “asparagine”, “aspartic acid”, “cysteine”, “glutamic acid”, “glutamine”, “glycine”, “histidine”, “isoleucine”, “leucine”, “lysine”, “methionine”, “phenylalanine”, “proline”, “serine”, “threonine”, “tryptophan”, “tyrosine”, “valine”, “ala”, “arg”, “asn”, “asp”, “cys”, “gln”, “glu”, “gly”, “his”, “ile”, “leu”, “lys”, “met”, “phe”, “pro”, “ser”, “thr”, “trp”, “tyr”, “val”

**Meaningful fuzzy matches**

“harpin”, “lysin”

### Note S5. Function Summarization Prompt User Prompt

You are a molecular and cell biology expert. Given this list of protein names from a single **bacterial** protein cluster, provide a one-sentence molecular or biological function description of this cluster in less than 35 words.

Protein names:

{name_list}

**Critical**

- NEVER invent functions for unrecognized proteins.
- Focus on the SHARED function across these proteins. If the proteins serve multiple unrelated functions, list them separately
- Use your training knowledge first. If confidence is low (*<*medium), use web search to verify against UniProt/NCBI databases or literature
- For Pfam domain identifiers (PF#####), use web search against InterPro/Pfam databases for any functional terms
- Be specific about substrates, cofactors, or pathways when mentioned
- If uncertain or ambiguous, indicate this clearly via confidence qualifiers (e.g., ’likely’, ’putative’, ’predicted’)
- If confidence is low, state: Functional details of this cluster are uncertain.

Do not describe your search, reasoning, or annotation process. Provide ONLY the functional description, nothing else.

**Parameters**

This prompt was used with an output token limit (*max_tokens*) of 150 but otherwise default parameters (*temperature* of 1.0, no extended thinking). The processed list of protein names for a Fusion function was put in place of *{name_list}* for each request.

### Note S6. Likelihood of Protein Name under Function Summary Prompt User Prompt

Given a protein with this function:

{raw_protein_name}

What is the numeric likelihood (0-10, with 10 as most likely) that it belongs to a cluster with this description:

{func_anno}

The output must contain only the numeric likelihood (0-10); nothing else is allowed under any circumstances.

**Parameters**

This prompt was used with an output token limit (*max_tokens*) of 10 but otherwise default parameters (*temperature* of 1.0, no extended thinking). A raw protein name was put in place of *{raw_protein_name}* and an LLM-generated function summary was put in place of *{func_anno}* for each request.

### Note S7. GO Extraction Prompt System Prompt

You are a cautious expert bioinformatics curator assigning Gene Ontology (GO) terms to proteins.

Goal:

Map protein descriptions to GO terms ONLY when the description clearly supports the annotation.

Output format (strict):

GO:XXXXXXX reliability=LEVEL; GO:XXXXXXX reliability=LEVEL or

” reliability=LOW

LEVEL must be: HIGH, MEDIUM, or LOW.

CRITICAL:

Output EXACTLY one line in this format.

Do not deliberate, reconsider, or revise in the output.

Stop immediately after producing the output line.

Examples:

DNA helicase -*>* GO:0004386 reliability=HIGH

Bacterial mobile genetic element transposases mediating DNA transposition and insertion sequence mobility through site-specific recombination and genome rearrangement mechanisms. -*>* GO:0006313 reliability=HIGH;

GO:0004803 reliability=HIGH; GO:0003677 reliability=MEDIUM mitochondrial ribosomal protein L3 -*>* GO:0005761 reliability=MEDIUM putative membrane protein -*>* ” reliability=LOW

Rules:

- Prefer precision over recall.
- Only output GO IDs that exist in the GO-basic ontology.
- Only assign GO terms directly supported by explicit functional or localization words in the description.
- Prefer the most specific GO term supported by the description.
- If evidence is weak or ambiguous, return ” reliability=LOW.
- Do NOT output broad root terms: GO:0008150, GO:0003674, GO:0005575.
- Ignore non-functional words such as organism names, isoform labels, “putative”, “probable”, “predicted”, and “uncharacterized.”
- Remove GO terms not directly supported by the description.

Reliability:

HIGH = explicit enzyme or clear function

MEDIUM = function implied but indirect

LOW = weak evidence or vague description

The output must contain only GO IDs and the reliability label; nothing else is allowed under any circumstances.

**User Prompt**

Protein description:

{cluster_anno}

Task:

Return GO terms.

**Parameters**

This prompt was used with an output token limit (*max_tokens*) of 150 and *temperature* of 0 but otherwise default parameters (no extended thinking). The LLM-generated function summary for a Fusion function was put in place of *{cluster_anno}* for each request. The *temperature* parameter was removed for Opus 5, and we instead set the *effort* parameter to *low*.

**Post-processing**

We extracted the last match in the output response matching the following regular expression:

(GO:\d{7} reliability=(HIGH|MEDIUM|LOW)(; GO:\d{7} reliability=(HIGH|MEDIUM|LOW))*|\’\’ reliability=LOW)

### Note S8. GO Evaluation Metrics

In comparing functional ground truth, i.e. Swiss-Prot-derived GO term annotations, vs. LLM text-extracted GO terms, we measured Jaccard similarity (Eq. 1) and Wang similarity (Eq. 2; [37]) as well as minimum semantic distance, F1, precision, and recall weighted by information accretion (IA) (Eqs. 3–4) as described in [38] for each of the three GO subontologies (molecular functions, MF, biological processes, BP, and cellular components, CC), individually.

Jaccard Similarity carries the GO-term overlap of each Fusion function’s LLM-derived (*G_p_*) vs. ground truth (*G_t_*) labels, each including all ancestors.

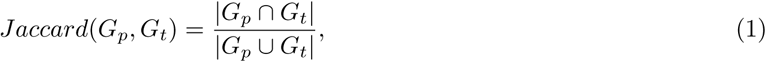

Wang similarity additionally incorporates GO’s ontology structure into its evaluations. *S*(*g, G^′^*) indicates semantic similarity of a term *g* to a set *G^′^*, where each term contributes a value that decays by a factor of 0.8 per “is_a” step from its descendants and 0.6 per “part_of” step from its descendants. The set-to-set score is the mean best-match over both directions,

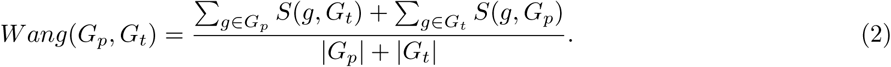

We computed both Jaccard and Wang similarity with *goatools* v1.5.2 [39]. For the information-accretion–weighted metrics, each term *g* carries an information accretion *ia*(*g*) value [38], provided in the CAFA 6 challenge [23] (precomputed from the June 1, 2025 *go-basic* graph). Weighted precision and recall reflect the shared information accretion over the predicted and true term sets

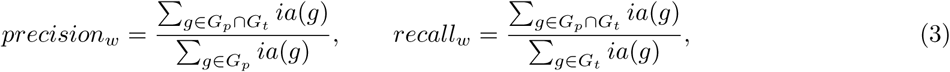

and weighted F1 is their harmonic mean,

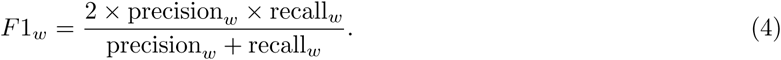

We computed these using the CAFA-evaluator-PK [40, 41] (-no_orphans to exclude root nodes, -norm pred) and our own derivative code.

All of the above are defined per protein or function, and we report each metric averaged over the *N_eval_* proteins (or functions) that have both a nonempty ground-truth and nonempty predicted GO sets across the compared methods. For the IA-weighted metrics, we reported *macro-averages*, where precision, recall, and F1 are calculated across all annotation of each protein or function and simply averaged, and *micro-averages*, where these metrics are computed across all (sample) weighted true positives, false positives, and false negatives.

All statistical testing was carried out using the ‘scipy‘ v1.15.3 Python package, specifically with scipy.stats.ttest_rel, scipy.stats.pearsonr, scipy.stats.spearmanr, and scipy.stats.ks_2samp.

---

**Figure S1:**
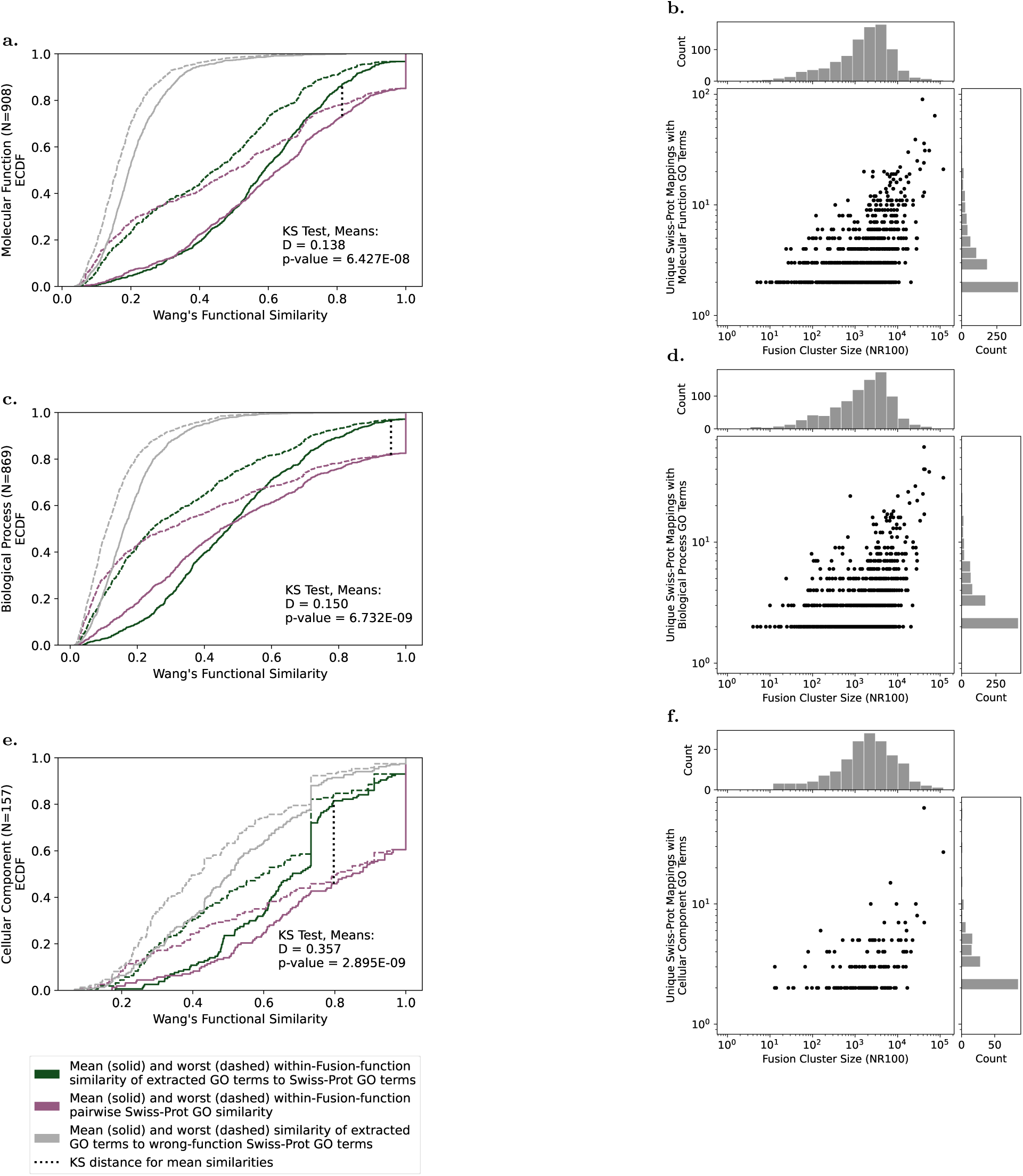
Variety of GO terms within a Fusion function bounds maximum accuracy of text-derived GO terms. Empirical cumulative distribution functions (ECDFs) of per Fusion function (1) within-function pairwise similarities of experimentally validated Swiss-Prot (SP) GO terms (purple) and (2) similarities between GO terms extracted by Opus 5 from Haiku-3.5-generated function summaries and SP GO terms for the corresponding Fusion function (dark green) and a randomly sampled other Fusion function (gray) are shown, and lower lines indicate higher accuracy (a,c,e). Numbers of unique SP mappings with GO terms roughly increase with cluster size (b,d,f). All Fusion functions with GO terms from at least two SP entries and at least one GO term extracted with Opus 5 from the Haiku-3.5-generated function summary per subontology are considered. Kolmogorov-Smirnov tests (at p-value cutoff of 0.05) indicate that the within function GO term variety vs. text-derived GO term variety distributions are significantly different, though they are noticeably closer to each other than the wrong-function baseline.

**Table S1:** GO terms extracted from text vs. Swiss-Prot annotations.

| Sub-ontology | Text* | $N_{gt}$ | $N_{int}$ | $N_{eval}$ | Weighted F1 | Weighted Precision | Weighted Recall | Wang Similarity |
| --- | --- | --- | --- | --- | --- | --- | --- | --- |
| Molecular Function | Protein Name | 657 | 595 | 464 | **0.78 (0.76-0.80)** | **0.89 (0.87-0.91)** | **0.75 (0.73-0.77)** | **0.79 (0.77-0.80)** |
|  | Curated Protein Function Text | 574 | 505 | 464 | **0.73 (0.71-0.75)** | 0.85 (0.83-0.87) | **0.71 (0.69-0.73)** | **0.74 (0.73-0.76)** |
|  | LLM Function-Cluster Summary | 657 | 604 | 464 | 0.67 (0.65-0.70) | 0.81 (0.79-0.83) | 0.65 (0.62-0.67) | 0.70 (0.68-0.71) |
| Biological Process | Protein Name | 558 | 477 | 400 | **0.73 (0.71-0.75)** | **0.86 (0.84-0.88)** | 0.69 (0.67-0.72) | **0.74 (0.72-0.76)** |
|  | Curated Protein Function Text | 502 | 455 | 400 | 0.72 (0.69-0.74) | 0.79 (0.77-0.82) | 0.72 (0.70-0.75) | 0.73 (0.71-0.75) |
|  | LLM Function-Cluster Summary | 558 | 539 | 400 | 0.68 (0.66-0.70) | 0.76 (0.74-0.79) | 0.69 (0.67-0.71) | 0.69 (0.67-0.71) |
| Cellular Component | Protein Name | 538 | 211 | 94 | 0.77 (0.72-0.82) | 0.93 (0.89-0.97) | 0.74 (0.68-0.79) | 0.74 (0.68-0.79) |
|  | Curated Protein Function Text | 451 | 144 | 94 | 0.74 (0.69-0.79) | 0.85 (0.79-0.90) | 0.75 (0.70-0.80) | 0.75 (0.70-0.80) |
|  | LLM Function-Cluster Summary | 538 | 203 | 94 | 0.72 (0.67-0.77) | 0.90 (0.85-0.94) | 0.68 (0.63-0.74) | 0.68 (0.63-0.74) |

\* We used Opus 5 to extract GO terms from (1) Swiss-Prot protein names (N=795), (2) Swiss-Prot protein function text (N=647), and (3) LLM (Haiku)-generated Fusion function-cluster summaries (N=795).

**Table S2:**
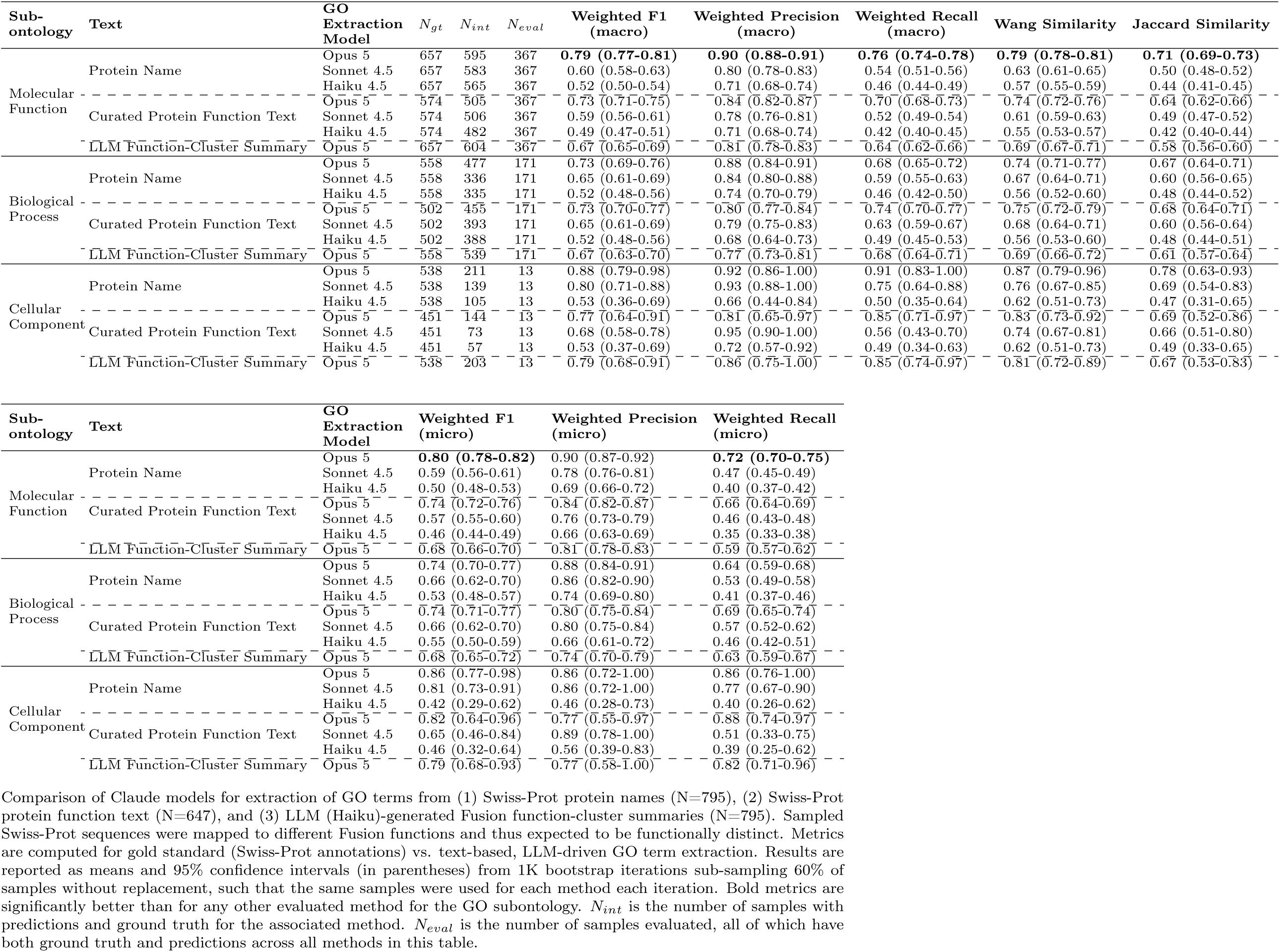
GO terms extracted from text vs. Swiss-Prot annotations.

**Table S3:** Correlation between LLM (Haiku 4.5) confidence in function summaries and accuracy of GO terms extracted from these summaries.

| Subontology | Similarity | Spearman | | Pearson | | $N_{int}$ |
| --- | --- | --- | --- | --- | --- | --- |
| | | $\rho$ | p | r | p | |
| Biological Process | Wang | 0.22 | 5.0e-10 | 0.18 | 3.6e-7 | 783 |
|  | Jaccard | 0.25 | 9.1e-13 | 0.21 | 3.7e-9 |  |
| Molecular Function | Wang | 0.04 | 0.26 | 0.08 | 0.02 | 761 |
|  | Jaccard | 0.09 | 0.02 | 0.09 | 0.01 |  |
| Cellular Component | Wang | 0.05 | 0.35 | -0.03 | 0.61 | 305 |
|  | Jaccard | 0.02 | 0.70 | -0.06 | 0.29 |  |

Haiku 4.5 was prompted to estimate the likelihood (0-10) of a protein name belonging to a Haiku-3.5-generated function summary. For each of $N = 1000$ function summaries, we averaged such likelihood estimates for 10 protein names sampled with replacement from the set of protein names used to generate the functional summary (via LLM prompting). Correlation was then done between the average LLM-estimated likelihood of protein names for each functional summary against the accuracy of the GO terms extracted with Opus 4.6 from the functional summary, with the ground truth as the GO terms mapped by homology to Swiss-Prot for the corresponding Fusion function (*Fusion-SP*). $N_{int}$ is the number of samples with both *Fusion-SP* and Opus-extracted GO terms for the sub-ontology, which is equivalent to the number of samples for which we performed the correlation tests.

**Table S4:** Predicting CAFA Molecular Function (MF) GO terms.

| Method | $N_{int}$ | $N_{eval}$ | Weighted F1 | Weighted Precision | Weighted Recall | Wang Similarity |
| --- | --- | --- | --- | --- | --- | --- |
| Best SP | 496 | 394 | 0.61 (0.59-0.64) | 0.63 (0.61-0.66) | 0.73 (0.70-0.75) | 0.64 (0.62-0.66) |
| All SP | 502 | 394 | 0.61 (0.58-0.63) | 0.59 (0.56-0.62) | 0.78 (0.76-0.80) | 0.65 (0.63-0.66) |
| Fusion-SP | 627 | 394 | 0.57 (0.55-0.60) | 0.51 (0.49-0.54) | **0.85 (0.83-0.87)** | 0.64 (0.62-0.66) |
| Opus 4.6 Text, Opus 4.6 GO | 632 | 394 | 0.51 (0.48-0.54) | 0.56 (0.54-0.59) | 0.57 (0.54-0.60) | 0.55 (0.53-0.57) |
| Haiku 3.5 Text, Opus 5 GO | 593 | 394 | 0.54 (0.51-0.57) | 0.61 (0.58-0.64) | 0.58 (0.55-0.61) | 0.57 (0.55-0.59) |
| Haiku 3.5 Text, Opus 4.6 GO | 595 | 394 | 0.49 (0.46-0.51) | 0.57 (0.54-0.59) | 0.51 (0.48-0.54) | 0.52 (0.51-0.54) |
| Haiku 3.5 Text, Haiku 4.5 GO | 565 | 394 | 0.38 (0.35-0.40) | 0.53 (0.50-0.56) | 0.37 (0.34-0.39) | 0.45 (0.43-0.47) |

*Model-selection dataset* proteins were assigned GO terms using homology-based and LLM-driven methods. Annotations were compared to CAFA 6 Protein Function Prediction Challenge [23] -provided GO terms. *Haiku 3.5 Text*, *Opus 5 GO* (underlined) was our selected model combination and what is provided in the web applications. Results are reported as macro-averaged means and 95% confidence intervals (in parentheses) from 1K bootstrap iterations sub-sampling 60% of samples without replacement, such that the same samples were used for each method each iteration. Of all metrics, only weighted recall was significantly better (bold) for one method (*Fusion-SP*) than all other compared methods. $N_{int}$ is the number of samples with predictions and ground truth for the associated method. $N_{eval}$ is the number of samples evaluated, all of which have both ground truth and predictions across all methods in this table. $N_{gt} = 691$ is the number of proteins in the sample with ground truth MF GO terms. Tables S5 and S6 contain results for additional metrics and GO subontologies.

**Table S5:** Annotating CAFA 6 Protein Function Prediction Challenge [23] GO terms by identifying GO-term annotated Swiss-Prot homologs.

| Subontology | Method | $N_{gt}$ | $N_{int}$ | $N_{eval}$ | Weighted F1<br>(macro) | Weighted Precision<br>(macro) | Weighted Recall<br>(macro) | Wang Similarity | Jaccard Similarity |
| --- | --- | --- | --- | --- | --- | --- | --- | --- | --- |
| Molecular<br>Function | Best SP | 691 | 496 | 495 | 0.62 (0.59-0.64) | 0.64 (0.61-0.67) | 0.72 (0.70-0.74) | 0.64 (0.62-0.66) | 0.54 (0.52-0.57) |
|  | All SP | 691 | 502 | 495 | 0.61 (0.59-0.63) | 0.60 (0.57-0.62) | 0.77 (0.75-0.80) | 0.65 (0.63-0.67) | 0.54 (0.52-0.56) |
|  | Fusion-SP | 691 | 627 | 495 | 0.59 (0.56-0.61) | 0.53 (0.51-0.56) | **0.86 (0.84-0.88)** | 0.65 (0.63-0.67) | 0.52 (0.50-0.54) |
| Biological<br>Process | Best SP | 640 | 401 | 401 | 0.63 (0.60-0.65) | 0.70 (0.67-0.73) | 0.65 (0.62-0.68) | 0.64 (0.62-0.66) | 0.57 (0.54-0.59) |
|  | All SP | 640 | 435 | 401 | 0.60 (0.58-0.63) | 0.64 (0.61-0.67) | 0.68 (0.65-0.71) | 0.63 (0.61-0.66) | 0.55 (0.52-0.58) |
|  | Fusion-SP | 640 | 587 | 401 | 0.62 (0.60-0.65) | 0.58 (0.56-0.61) | **0.84 (0.81-0.86)** | 0.66 (0.64-0.68) | 0.56 (0.53-0.58) |
| Cellular<br>Component | Best SP | 567 | 366 | 366 | 0.61 (0.58-0.64) | 0.76 (0.73-0.79) | 0.60 (0.57-0.64) | 0.76 (0.74-0.78) | 0.63 (0.61-0.66) |
|  | All SP | 567 | 390 | 366 | 0.61 (0.58-0.64) | 0.69 (0.66-0.73) | 0.67 (0.63-0.70) | 0.75 (0.73-0.77) | 0.60 (0.58-0.63) |
|  | Fusion-SP | 567 | 517 | 366 | 0.60 (0.57-0.63) | 0.63 (0.60-0.66) | **0.75 (0.72-0.78)** | 0.75 (0.73-0.77) | 0.58 (0.55-0.61) |

| Subontology | Method | Weighted F1<br>(micro) | Weighted Precision<br>(micro) | Weighted Recall<br>(micro) |
| --- | --- | --- | --- | --- |
| Molecular<br>Function | Best SP | 0.66 (0.63-0.67) | **0.60 (0.57-0.63)** | 0.72 (0.69-0.74) |
|  | All SP | 0.62 (0.60-0.65) | 0.52 (0.49-0.55) | 0.77 (0.75-0.80) |
|  | Fusion-SP | 0.53 (0.50-0.57) | 0.39 (0.35-0.43) | **0.86 (0.84-0.89)** |
| Biological<br>Process | Best SP | **0.65 (0.62-0.67)** | **0.68 (0.65-0.71)** | 0.62 (0.59-0.65) |
|  | All SP | 0.54 (0.50-0.59) | 0.46 (0.41-0.52) | 0.67 (0.63-0.70) |
|  | Fusion-SP | 0.48 (0.43-0.54) | 0.34 (0.29-0.40) | **0.82 (0.80-0.85)** |
| Cellular<br>Component | Best SP | 0.60 (0.56-0.64) | **0.68 (0.63-0.74)** | 0.54 (0.49-0.58) |
|  | All SP | 0.50 (0.45-0.57) | 0.42 (0.36-0.51) | 0.63 (0.59-0.66) |
|  | Fusion-SP | 0.50 (0.46-0.56) | 0.38 (0.33-0.45) | **0.73 (0.69-0.77)** |

*Model-selection dataset* proteins ( $N = 1000$ ) were assigned GO terms using homology based methods. Proteins were aligned directly to Swiss-Prot and GO terms were transferred from the Swiss-Prot protein (excluding self-hits) with highest $HFS \geq 14$ (*Best SP*) or all non-self-hit Swiss-Prot matches with $HFS \geq 14$ (*All SP*). Alternatively, proteins were mapped to a Fusion function and then the GO terms mapped from Swiss-Prot to the Fusion function via alignment were assigned to the query protein (*Fusion-SP*). The ground truth were the CAFA 6 Protein Function Prediction Challenge [23] -provided GO terms. Results are reported as means and 95% confidence intervals (in parentheses) from 1K bootstrap iterations sub-sampling 60% of samples without replacement, such that the same samples were used for each method each iteration. Bold metrics are significantly better than for any other evaluated method for the GO subontology. $N_{int}$ is the number of samples with predictions and ground truth for the associated method. $N_{eval}$ is the number of samples evaluated, all of which have both ground truth and predictions across all methods in this table.

**Table S6:** Annotating CAFA 6 Protein Function Prediction Challenge [23] GO terms by extraction from LLM-generated Fusion function summaries.

| Subontology | Method | $N_{gt}$ | $N_{int}$ | $N_{eval}$ | Weighted F1<br>(macro) | Weighted Precision<br>(macro) | Weighted Recall<br>(macro) | Wang<br>Similarity | Jaccard<br>Similarity |
| --- | --- | --- | --- | --- | --- | --- | --- | --- | --- |
| Molecular<br>Function | Opus 4.6 Only | 691 | 632 | 472 | 0.50 (0.47-0.52) | 0.55 (0.52-0.57) | 0.56 (0.54-0.59) | 0.55 (0.53-0.57) | 0.44 (0.43-0.46) |
|  | Haiku 3.5 Text, Opus 5 GO | 691 | 593 | 472 | 0.52 (0.50-0.54) | 0.59 (0.56-0.62) | 0.57 (0.54-0.59) | 0.57 (0.55-0.58) | 0.47 (0.45-0.49) |
|  | Haiku 3.5 Text, Opus 4.6 GO | 691 | 595 | 472 | 0.48 (0.46-0.51) | 0.56 (0.53-0.59) | 0.51 (0.49-0.53) | 0.53 (0.51-0.55) | 0.43 (0.41-0.45) |
|  | Sonnet 4.5 Only | 691 | 611 | 472 | 0.47 (0.44-0.49) | 0.56 (0.53-0.59) | 0.49 (0.46-0.52) | 0.52 (0.51-0.54) | 0.42 (0.40-0.44) |
|  | Haiku 3.5 Text, Sonnet 4.5 GO | 691 | 567 | 472 | 0.46 (0.44-0.49) | 0.59 (0.56-0.62) | 0.46 (0.44-0.49) | 0.52 (0.50-0.54) | 0.42 (0.40-0.44) |
|  | Haiku 3.5 Text, Haiku 4.5 GO | 691 | 565 | 472 | 0.38 (0.36-0.40) | 0.53 (0.50-0.56) | 0.38 (0.35-0.40) | 0.46 (0.45-0.48) | 0.36 (0.34-0.38) |
| Biological<br>Process | Opus 4.6 Only | 640 | 604 | 425 | 0.51 (0.48-0.54) | 0.60 (0.57-0.63) | 0.52 (0.49-0.55) | 0.55 (0.53-0.58) | 0.46 (0.44-0.49) |
|  | Haiku 3.5 Text, Opus 5 GO | 640 | 597 | 425 | 0.52 (0.49-0.54) | 0.62 (0.59-0.65) | 0.53 (0.50-0.56) | 0.56 (0.53-0.58) | 0.46 (0.44-0.49) |
|  | Haiku 3.5 Text, Opus 4.6 GO | 640 | 610 | 425 | 0.49 (0.47-0.52) | 0.59 (0.56-0.62) | 0.50 (0.48-0.53) | 0.54 (0.52-0.56) | 0.45 (0.42-0.47) |
|  | Sonnet 4.5 Only | 640 | 521 | 425 | 0.49 (0.46-0.52) | 0.62 (0.59-0.65) | 0.48 (0.45-0.51) | 0.54 (0.51-0.56) | 0.45 (0.43-0.48) |
|  | Haiku 3.5 Text, Sonnet 4.5 GO | 640 | 568 | 425 | 0.47 (0.44-0.49) | 0.60 (0.57-0.63) | 0.46 (0.43-0.48) | 0.51 (0.49-0.53) | 0.43 (0.40-0.45) |
|  | Haiku 3.5 Text, Haiku 4.5 GO | 640 | 557 | 425 | 0.39 (0.37-0.42) | 0.54 (0.51-0.56) | 0.38 (0.36-0.41) | 0.46 (0.44-0.48) | 0.36 (0.34-0.38) |
| Cellular<br>Component | Opus 4.6 Only | 567 | 270 | 66 | 0.59 (0.51-0.66) | 0.66 (0.57-0.75) | 0.60 (0.51-0.68) | 0.69 (0.63-0.74) | 0.58 (0.51-0.64) |
|  | Haiku 3.5 Text, Opus 5 GO | 567 | 192 | 66 | 0.63 (0.56-0.70) | 0.75 (0.68-0.83) | 0.63 (0.55-0.71) | 0.70 (0.66-0.75) | 0.62 (0.56-0.68) |
|  | Haiku 3.5 Text, Opus 4.6 GO | 567 | 233 | 66 | 0.65 (0.57-0.72) | 0.77 (0.69-0.85) | 0.63 (0.56-0.71) | 0.72 (0.66-0.76) | 0.63 (0.56-0.69) |
|  | Sonnet 4.5 Only | 567 | 186 | 66 | 0.56 (0.48-0.62) | 0.73 (0.66-0.81) | 0.53 (0.45-0.60) | 0.66 (0.62-0.71) | 0.57 (0.51-0.63) |
|  | Haiku 3.5 Text, Sonnet 4.5 GO | 567 | 162 | 66 | 0.53 (0.47-0.60) | 0.72 (0.64-0.81) | 0.50 (0.43-0.57) | 0.64 (0.60-0.69) | 0.55 (0.49-0.61) |
|  | Haiku 3.5 Text, Haiku 4.5 GO | 567 | 113 | 66 | 0.47 (0.40-0.55) | 0.64 (0.55-0.72) | 0.44 (0.36-0.51) | 0.61 (0.56-0.66) | 0.49 (0.42-0.56) |

| Subontology | Method | Weighted F1<br>(micro) | Weighted Precision<br>(micro) | Weighted Recall<br>(micro) |
| --- | --- | --- | --- | --- |
| Molecular<br>Function | Opus 4.6 Only | 0.53 (0.51-0.55) | 0.51 (0.48-0.53) | 0.55 (0.52-0.57) |
|  | Haiku 3.5 Text, Opus 5 GO | 0.55 (0.53-0.57) | 0.56 (0.53-0.59) | 0.54 (0.52-0.57) |
|  | Haiku 3.5 Text, Opus 4.6 GO | 0.51 (0.49-0.53) | 0.53 (0.50-0.56) | 0.49 (0.47-0.52) |
|  | Sonnet 4.5 Only | 0.49 (0.47-0.51) | 0.52 (0.49-0.55) | 0.47 (0.44-0.49) |
|  | Haiku 3.5 Text, Sonnet 4.5 GO | 0.49 (0.47-0.51) | 0.56 (0.53-0.59) | 0.44 (0.42-0.46) |
|  | Haiku 3.5 Text, Haiku 4.5 GO | 0.39 (0.36-0.41) | 0.46 (0.43-0.49) | 0.33 (0.31-0.36) |
| Biological<br>Process | Opus 4.6 Only | 0.52 (0.49-0.55) | 0.57 (0.54-0.60) | 0.47 (0.45-0.50) |
|  | Haiku 3.5 Text, Opus 5 GO | 0.53 (0.50-0.55) | 0.58 (0.56-0.61) | 0.48 (0.45-0.51) |
|  | Haiku 3.5 Text, Opus 4.6 GO | 0.50 (0.48-0.53) | 0.56 (0.53-0.59) | 0.46 (0.43-0.48) |
|  | Sonnet 4.5 Only | 0.50 (0.48-0.53) | 0.59 (0.56-0.63) | 0.44 (0.41-0.47) |
|  | Haiku 3.5 Text, Sonnet 4.5 GO | 0.47 (0.45-0.50) | 0.57 (0.54-0.60) | 0.41 (0.38-0.44) |
|  | Haiku 3.5 Text, Haiku 4.5 GO | 0.40 (0.38-0.42) | 0.50 (0.47-0.53) | 0.33 (0.31-0.36) |
| Cellular<br>Component | Opus 4.6 Only | 0.57 (0.48-0.67) | 0.58 (0.48-0.68) | 0.57 (0.47-0.67) |
|  | Haiku 3.5 Text, Opus 5 GO | 0.69 (0.61-0.77) | 0.69 (0.60-0.80) | 0.69 (0.61-0.77) |
|  | Haiku 3.5 Text, Opus 4.6 GO | 0.68 (0.59-0.77) | 0.71 (0.62-0.81) | 0.65 (0.55-0.76) |
|  | Sonnet 4.5 Only | 0.60 (0.52-0.67) | 0.64 (0.54-0.73) | 0.57 (0.49-0.66) |
|  | Haiku 3.5 Text, Sonnet 4.5 GO | 0.59 (0.50-0.67) | 0.62 (0.52-0.71) | 0.57 (0.47-0.67) |
|  | Haiku 3.5 Text, Haiku 4.5 GO | 0.44 (0.36-0.53) | 0.55 (0.45-0.65) | 0.37 (0.29-0.46) |

*Model-selection dataset* proteins ( $N = 1000$ ) were assigned GO extracted with an LLM from an LLM-generated function summary. The ground truth were the CAFA 6 Protein Function Prediction Challenge [23] -provided GO terms. *Haiku 3.5 Text*, *Opus 4.6 GO* refers to GO terms extracted using Opus 4.6 from functions summarized by Haiku 3.5, *Opus 4.6 Only* refers to GO terms extracted using Opus 4.6 from functions summarized by Opus 4.6, etc. *Haiku 3.5 Text*, *Opus 5 GO* (underlined) was our chosen method and what is provided in the web applications. Results are reported as means and 95% confidence intervals (in parentheses) from 1K bootstrap iterations sub-sampling 60% of samples without replacement, such that the same samples were used for each method each iteration. No metrics for one method are significantly better than for any other evaluated method for a given GO subontology. $N_{int}$ is the number of samples with predictions and ground truth for the associated method. $N_{eval}$ is the number of samples evaluated, all of which have both ground truth and predictions across all methods in this table.

**Table S7:** Evaluation of Opus 4.6 LLM-categorized GO reliability against CAFA 6 Protein Function Prediction Challenge [23] GO terms.

| Subontology | Reliability Levels | $N_{gt}$ | $N_{int}$ | $N_{eval}$ | Weighted F1<br>(macro) | Weighted Precision<br>(macro) | Weighted Recall<br>(macro) | Wang<br>Similarity | Jaccard<br>Similarity |
| --- | --- | --- | --- | --- | --- | --- | --- | --- | --- |
| Molecular<br>Function | High | 691 | 384 | 384 | 0.53 (0.50-0.55) | 0.68 (0.65-0.71) | 0.50 (0.47-0.53) | 0.55 (0.53-0.57) | 0.46 (0.44-0.49) |
|  | Medium and High | 691 | 584 | 384 | 0.56 (0.53-0.59) | 0.63 (0.60-0.66) | 0.60 (0.57-0.63) | 0.59 (0.56-0.61) | 0.49 (0.47-0.51) |
|  | Low, Medium, and High | 691 | 593 | 384 | 0.56 (0.53-0.59) | 0.63 (0.60-0.66) | 0.60 (0.58-0.63) | 0.59 (0.57-0.61) | 0.49 (0.47-0.52) |
| Biological<br>Process | High | 640 | 359 | 359 | 0.55 (0.52-0.58) | 0.73 (0.70-0.77) | 0.49 (0.47-0.52) | 0.58 (0.56-0.61) | 0.51 (0.48-0.54) |
|  | Medium and High | 640 | 571 | 359 | 0.55 (0.53-0.58) | 0.67 (0.64-0.70) | 0.56 (0.53-0.59) | 0.60 (0.57-0.62) | 0.51 (0.48-0.53) |
|  | Low, Medium, and High | 640 | 597 | 359 | 0.55 (0.53-0.58) | 0.66 (0.63-0.69) | 0.56 (0.54-0.59) | 0.59 (0.57-0.62) | 0.50 (0.48-0.53) |
| Cellular<br>Component | High | 567 | 40 | 40 | 0.57 (0.47-0.67) | 0.70 (0.60-0.81) | 0.54 (0.45-0.64) | 0.64 (0.58-0.71) | 0.54 (0.46-0.63) |
|  | Medium and High | 567 | 162 | 40 | 0.61 (0.52-0.71) | 0.68 (0.57-0.79) | 0.62 (0.52-0.72) | 0.67 (0.60-0.74) | 0.57 (0.48-0.66) |
|  | Low, Medium, and High | 567 | 192 | 40 | 0.62 (0.52-0.71) | 0.68 (0.57-0.78) | 0.64 (0.54-0.73) | 0.67 (0.60-0.74) | 0.57 (0.48-0.65) |

| Subontology | Reliability Levels | Weighted F1<br>(micro) | Weighted Precision<br>(micro) | Weighted Recall<br>(micro) |
| --- | --- | --- | --- | --- |
| Molecular<br>Function | High | 0.55 (0.53-0.58) | 0.66 (0.63-0.70) | 0.48 (0.45-0.50) |
|  | Medium and High | 0.59 (0.56-0.61) | 0.59 (0.56-0.63) | 0.58 (0.55-0.61) |
|  | Low, Medium, and High | 0.59 (0.56-0.61) | 0.59 (0.56-0.63) | 0.58 (0.55-0.61) |
| Biological<br>Process | High | 0.56 (0.53-0.59) | **0.75 (0.71-0.78)** | 0.45 (0.42-0.48) |
|  | Medium and High | 0.56 (0.53-0.59) | 0.63 (0.60-0.67) | 0.50 (0.47-0.53) |
|  | Low, Medium, and High | 0.56 (0.53-0.59) | 0.63 (0.60-0.66) | 0.51 (0.48-0.54) |
| Cellular<br>Component | High | 0.63 (0.51-0.74) | 0.67 (0.53-0.79) | 0.59 (0.48-0.72) |
|  | Medium and High | 0.68 (0.57-0.78) | 0.68 (0.55-0.79) | 0.68 (0.57-0.80) |
|  | Low, Medium, and High | 0.68 (0.58-0.78) | 0.67 (0.55-0.78) | 0.70 (0.59-0.81) |

Opus 5 was prompted to extract GO terms from Haiku-3.5-generated Fusion function summaries for the 1000 proteins in the *model-selection dataset* and categorize them as low-, medium-, or high-reliability. Here, we compare keeping GO terms across all reliability levels against keeping only high-reliability or high- and medium-reliability GO terms. The ground truth were the CAFA 6 Protein Function Prediction Challenge [23] -provided GO terms. Results are reported as means and 95% confidence intervals (in parentheses) from 1K bootstrap iterations sub-sampling 60% of samples without replacement, such that the same samples were used for each method each iteration. Bold metrics are significantly better than for the other reliability cutoffs for the GO subontology. $N_{int}$ is the number of samples with predictions and ground truth for the associated method. $N_{eval}$ is the number of samples evaluated and is equivalent to the number of proteins with high-reliability predicted GO terms.

**Table S8:** Performance on full CAFA 6 bacterial training set.

| Subontology | $N_{gt}$ | $N_{int}$ | Prediction Normalization (-norm pred) | | | CAFA Normalization (-norm cafa) | | | Weighted<br>F1 (micro) | Weighted<br>Precision<br>(micro) | Weighted<br>Recall<br>(micro) |
| --- | --- | --- | --- | --- | --- | --- | --- | --- | --- | --- | --- |
|  |  |  | Weighted<br>F1 (macro) | Weighted<br>Precision<br>(macro) | Weighted<br>Recall<br>(macro) | Weighted<br>F1 (macro) | Weighted<br>Precision<br>(macro) | Weighted<br>Recall<br>(macro) |  |  |  |
| Molecular Function | 4,199 | 3747 | 0.56 | 0.57 | 0.55 | 0.53 | 0.57 | 0.49 | 0.52 | 0.53 | 0.5 |
| Biological Process | 4,064 | 3717 | 0.53 | 0.58 | 0.49 | 0.51 | 0.58 | 0.45 | 0.48 | 0.55 | 0.42 |
| Cellular Component | 3,690 | 1265 | 0.61 | 0.8 | 0.49 | 0.28 | 0.8 | 0.17 | 0.3 | 0.67 | 0.19 |

Performance of LLM-extracted Fusion GO terms for all bacterial samples in the CAFA 6 Protein Function Prediction challenge [23]. 6,225 of the 6,251 total bacterial sequences were mapped to 3,897 unique Fusion clusters, and 6,183 of the 6,251 could be annotated with *highly meaningful* text. GO terms were extracted with Opus 5 from the *highly meaningful* text annotations, which were either Haiku-3.5-generated function summaries or single, *highly meaningful* protein names. We report macro-averaged metrics with both "prediction normalization," where recall is normalized by the number of samples with both ground truth and predictions for the subontology ( $N_{int}$ ), and with "CAFA normalization," where recall is normalized by the number of samples with ground truth for the subontology ( $N_{gt}$ ).

**Table S9:** Fusion expands metagenome/microbiome annotation. with *Highly*.

| SRA ID | $N_{qreads}$ | % Mapped to<br>EC Numbers<br>with *mifaser* | % Mapped to<br>Fusion Functions<br>with *Highly*<br>*Meaningful* Text |
| --- | --- | --- | --- |
| SRR1566021 | 4.1 M | 5.51% | 41.93% |
| SRR1569462 | 10.1 M | 5.36% | 40.18% |
| SRR1569742 | 40.3 M | 2.33% | 24.85% |
| SRR1569812 | 39.9 M | 2.20% | 23.98% |
| SRR1570801 | 41.5 M | 1.22% | 13.22% |
| SRR1570802 | 38.9 M | 1.17% | 12.87% |
| **Mean** |  | **2.96%** | **26.17%** |

Nearly ten-fold the amount of metagenomic reads that previously could be annotated with EC numbers using *mifaser* are now mapped to Fusion functions with informative text annotations. EC numbers were assigned using the *mifaser* web application (accessed August 26, 2026) with the *GS-24-all* reference database and quality control on, and Fusion functions were assigned with the same *faser* algorithm against the NR60 Fusion reference with singletons. Singletons make up less than 0.5% of matches across all samples. $N_{qreads}$ is the number of reads that passed quality control (*fastp* v0.20.1 [33] with *phredquality* = 20 and *readlength* = 40). Metagenomes are Deepwater Horizon oil spill beach sand samples from the pre-oil (SRR1566021, SRR1569462), oiled (SRR1569742, SRR1569812), and post-oil recovered (SRR1570801, SRR1570802) phases (NCBI BioProject PRJNA260285) [32].

